# Split-TurboID enables contact-dependent proximity labeling in cells

**DOI:** 10.1101/2020.03.11.988022

**Authors:** Kelvin F. Cho, Tess C. Branon, Sanjana Rajeev, Tanya Svinkina, Namrata D. Udeshi, Themis Thoudam, Chulhwan Kwak, Hyun-Woo Rhee, In-Kyu Lee, Steven A. Carr, Alice Y. Ting

## Abstract

Proximity labeling (PL) catalyzed by promiscuous enzymes such as TurboID have enabled the proteomic analysis of subcellular regions difficult or impossible to access by conventional fractionation-based approaches. Yet some cellular regions, such as organelle contact sites, remain out of reach for current PL methods. To address this limitation, we split the enzyme TurboID into two inactive fragments that recombine when driven together by a protein-protein interaction or membrane-membrane apposition. At endoplasmic reticulum (ER)-mitochondria contact sites, reconstituted TurboID catalyzed spatially-restricted biotinylation, enabling the enrichment and identification of >100 endogenous proteins, including many not previously linked to ER-mitochondria contacts. We validated eight novel candidates by biochemical fractionation and overexpression imaging. Overall, split-TurboID is a versatile tool for conditional and spatially-specific proximity labeling in cells.

## Introduction

Proximity labeling (PL) has proven to be a valuable tool for studying protein localization and interactions in living cells (1–3). In PL, a promiscuous enzyme such as APEX (4, 5), BioID (6), or TurboID (7) is genetically targeted to an organelle or protein complex of interest. Addition of a biotin-derived small molecule substrate then initiates biotinylation of endogenous proteins within a few nanometers of the promiscuous enzyme, via a diffusible radical intermediate in the case of APEX, or an activated biotin adenylate ester in the case of BioID and TurboID. After cell lysis, biotinylated proteins are harvested using streptavidin beads and identified by mass spectrometry.

Proximity labeling has been applied in many cell types and species to map the proteome composition of various organelles, including mitochondria (5, 8–10), synapses (11, 12), stress granules (13), and primary cilia (14). However, to increase the versatility of PL, new enzyme variants are needed. In particular, split enzymes could enable greater spatial specificity in the targeting of biotinylation activity, as well as PL activity that is conditional on a specific input, such as drug, calcium, or cell-cell contact. For example, contact sites between mitochondria and endoplasmic reticulum (ER) mediate diverse biology, from lipid biosynthesis to Ca^+2^ signaling to regulation of mitochondrial fission (15). There is great interest in probing the proteomic composition of ER-mitochondria contacts. However, direct fusion of a PL enzyme to one of the known ER-mitochondria contact resident proteins (e.g. Drp1 or Mff) would generate PL activity outside of ER-mitochondria contacts as well, because these proteins also reside in other subcellular locations (16, 17). On the other hand, use of a split PL enzyme, with one fragment targeted to the mitochondria, and the other targeted to the ER, would restrict biotinylation activity to ER-mitochondria contact sites specifically.

Split forms of APEX (18) and BioID (19–21) have previously been reported. However, split-APEX (developed by us) has not been used for proteomics, and the requirement for exogenous H_2_O_2_ and heme addition limits its utility in vivo. Split-BioID was first reported by De Munter et al. (19), followed by more active versions from Schopp et al. (20) and Kwak et al. (in press; (21)). All are derived from the parental enzyme BioID, which requires 18-24 hours of biotin labeling. We show below that the Schopp et al. split-BioID (20) does not exhibit activity in our hands, while the Kwak et al. split-BioID (21) requires 16+ hours of labeling to produce sufficient signal.

Hence we sought to develop an improved, more active split PL enzyme by starting from TurboID. In contrast to APEX, TurboID does not require any co-factors or co-oxidants; just biotin addition initiates labeling in cells or animals. TurboID is also >100-fold faster than BioID, requiring only 1-10 minutes of labeling time (7). We performed a screen of 14 different TurboID split sites to identify optimal fragments for high-affinity and low-affinity reconstitution. TurboID split at L73/G74 gave rapamycin-dependent reconstitution when fused to FRB and FKBP in multiple subcellular organelles. We then used split-TurboID to perform proteomic mapping of ER-mitochondria contact sites in mammalian cells. The resulting proteome of 101 proteins is highly specific and identifies many new mitochondria-ER contact site candidates, eight of which we validated by biochemical fractionation or overexpression imaging.

## Results

### Development of a split promiscuous biotin ligase with high activity

We started with TurboID, for the reasons given above, and sought to design split protein fragments with no detectable activity on their own, but high reconstituted activity. Given the diversity of ways in which split proteins are used, we envisioned engineering both a low-affinity fragment pair, whose reconstitution could be driven by a protein-protein or membrane-membrane association, and a high affinity pair that would reconstitute efficiently upon co-compartmentalization of fragments (Figure 1A). Previously, we developed split enzymes (split-APEX (18) and split-HRP (22)) by manually selecting cut sites in exposed loops, guided by protein crystal structures. Here, we instead utilized a recently-developed computational algorithm for predicting optimal protein split sites (23). SPELL (Split Protein rEassembly by Ligand or Light) calculates the energy profile of each candidate fragment relative to that of the full-length protein, and combines this information with solvent accessibility, sequence conservation, and geometric constraints to evaluate potential split sites, aiming for fragment pairs that give high reconstitution efficiency and minimal spontaneous assembly (23). Because crystal structures for TurboID and BioID are not available, we applied the SPELL algorithm to wild-type *E. coli* biotin ligase (BirA; PDB: 1HXD (24) and PDB: 2EWN (25)), from which both enzymes are derived.

**Figure 1.**
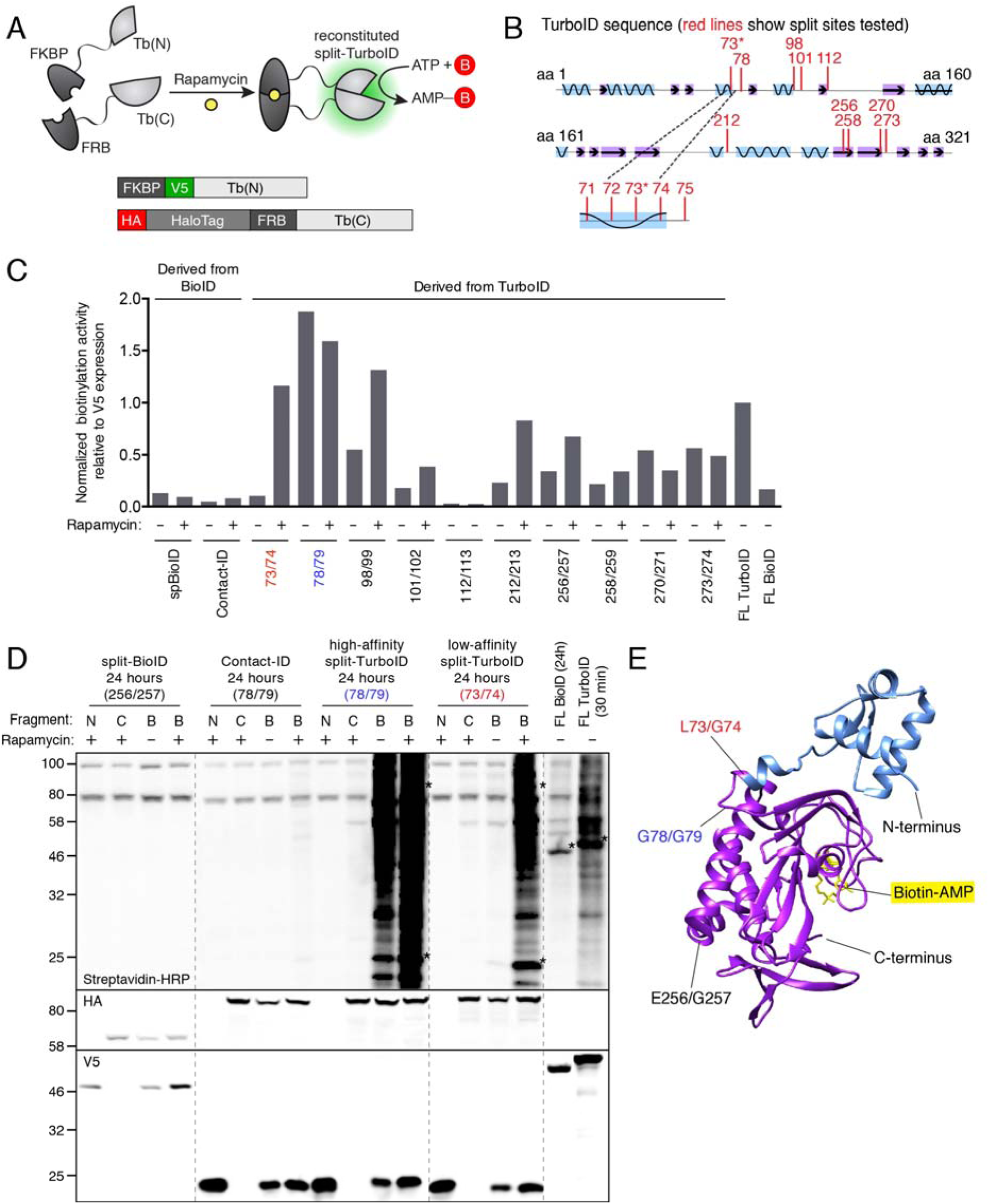
Engineering split-TurboID. *(A)* Schematic of split-TurboID reconstitution using the chemically-inducible FRB-FKBP dimerization system and design of constructs used to screen pairs of split sites. Upon rapamycin treatment, two inactive fragments of TurboID reconstitute to form an active enzyme capable of generating biotin-5’-AMP for promiscuous proximity-dependent labeling. N-terminal fragments (Tb(N)) were fused to FKBP and V5. C-terminal fragments (Tb(C)) were fused to HA, HaloTag, and FRB. The HaloTag was retained for initial screening as previous studies have shown that it can improve stability (18). *(B)* Split sites tested. Ten split sites were tested in the first round, with the 73/74 split being the best. In the second round, four additional sites were tested. Split sites are indicated as red lines along the TurboID protein sequence. The α helices are shown in blue and the β sheets are shown in purple. *(C)* Results of split site screen. Split-BioID (split at E256/G257) (20) and Contact-ID (split at G78/G79) (21) are previously described versions of split-BioID. Each fragment pair was tested in HEK293T cells with 24 hours biotin incubation in the presence or absence of rapamycin. At right, cells expressing full-length (FL) TurboID were incubated with biotin for 30 minutes. FL BioID was incubated with biotin for 24 hours. Cell lysates were analyzed by streptavidin blotting as in (D), and quantification was performed by dividing the streptavidin sum intensity by the anti-V5 intensity. Values were normalized to that of full-length TurboID. *(D)* Streptavidin blot comparing our final split-TurboID pair to full-length TurboID and BioID, and the previously-described split-BioID and Contact-ID pairs (20, 21). Labeling conditions were same as in (D). For each construct pair, lanes are shown with both fragments present (B), N-terminal fragment only (N), or C-terminal fragment only (C). Anti-V5 and anti-HA blotting show expression levels of N-terminal fragments (V5-tagged), C-terminal fragments (HA-tagged), and full-length enzymes (V5-tagged). Dashed lines indicate separate blots performed at the same time and developed simultaneously. Asterisks indicate ligase self-biotinylation. Full blots are shown in Supplementary Figure 1. *(E)* N- and C-terminal fragments (blue and purple, respectively) of split-TurboID (73/74), indicated on a structure of *E. coli* biotin ligase (PDB: 2EWN), from which TurboID was evolved (7). Biotin-AMP in the active site is shown in yellow. The low-affinity split-TurboID cut site is shown in red, the high-affinity split-TurboID cut site is shown in blue, and the previous split-BioID cut site is shown in black.

SPELL identified 10 potential split sites, all of which are in exposed loops. We rejected some of them based on prior experimental data: for example, cut site 62/63 was predicted by SPELL, but our previously-developed miniTurbo is truncated at amino acid 64 and retains high activity (7). We selected five of the SPELL-predicted cut sites for experimental testing (Figure 1B). In addition, we included in our screen five more cut sites used in previous split-BioIDs (20, 21). Each fragment pair was cloned as fusions to FKBP and FRB, proteins whose association can be induced by the small molecule rapamycin (Figure 1A). The constructs were all expressed in the cytosol of HEK293T cells and incubated with biotin for 24 hours in the presence or absence of rapamycin. Cell lysates were run on SDS-PAGE and blotted with streptavidin to evaluate the extent of promiscuous biotinylation. Figures 1C, D and Supplementary Figure 1A show that split-TurboIDs cut at 73/74, 78/79, and 98/99 give high reconstituted activity. 78/79 is the most active, under both +rapamycin and -rapamycin conditions, suggesting that the split fragments have high affinity for one another. The SPELL-predicted cut site 73/74 gave the greatest rapamycin-*dependent* activity, suggesting that it is a low-affinity, or conditional, split-TurboID.

We performed a secondary screen around the cut site 73/74 to further optimize low-affinity split-TurboID. Neighboring cut sites were tested (Supplementary Figure 1B), in addition to pairing of fragments with overlapping or gapped ends (Supplementary Figure 1C). None of these were better than the original 73/74 pair, so we selected this as our optimal low-affinity split-TurboID (referred to simply as “split-TurboID” in the remainder of the text).

In a side-by-side comparison to previous split-BioIDs (Figure 1C, D), both our high-affinity and low-affinity split-TurboIDs were far more active. The Kwak et al. split-BioID (also termed Contact ID (21)) showed rapamycin-dependent reconstitution with activity ∼12 fold lower than that of split-TurboID. This is consistent with the reported difference in catalytic activities of the parent enzymes TurboID and BioID (7). Interestingly, when the Contact-ID cut site (78/79) is used in TurboID, this yields our best high-affinity split-TurboID (78/79). The discrepancy between the rapamycin-dependence of Contact-ID and the rapamycin-*in*dependence of high-affinity split-TurboID is likely explained by their different regimes of activity; Contact-ID labeling may not be detectable in the -rapamaycin condition because the intrinsic activity is so low.

In our hands, the previously reported split-BioID from Schopp et al. (20) did not give any detectable signal over background after 24 hours of biotin incubation. Interestingly, TurboID split at the same position (256/257) did show some labeling (Figure 1C) but this activity was also observed with the N-terminal fragment alone (Supplementary Figure 1A), suggesting that this cut site may not yield a true protein complementation system. Notably, we found that the activity of split-TurboID is even greater than that of full-length BioID (Figure 1D; side-by-side comparison using 24 hours of biotin incubation), suggesting that split-TurboID’s activity level should be adequate for a wide range of applications.

By referencing the protein structure of *E. coli* biotin ligase (PDB: 2EWN), from which TurboID was evolved, we see that the split-TurboID site (L73/G74) separates the protein into two globular domains (Figure 1E). It is intriguing that just by moving the cut site five residues away (to 78/79), we produce a split-TurboID system that is high affinity/rapamycin-independent rather than low affinity/rapamycin-dependent.

### Further characterization of split-TurboID

To further characterize split-TurboID, we confirmed by confocal fluorescence imaging that the constructs catalyze biotinylation in a biotin- and rapamycin-dependent manner (Figure 2A). Reconstituted split-TurboID is not as active as full-length TurboID, but gave detectable biotinylation after just 30 minutes of biotin incubation (Figure 2B). To probe the kinetics of reconstitution, we compared rapamycin pre-incubation to rapamycin co-addition with biotin. There was no difference in biotinylation activity (Supplementary Figure 2A), suggesting that split-TurboID becomes active and begins catalyzing biotinylation immediately upon rapamycin addition.

**Figure 2.**
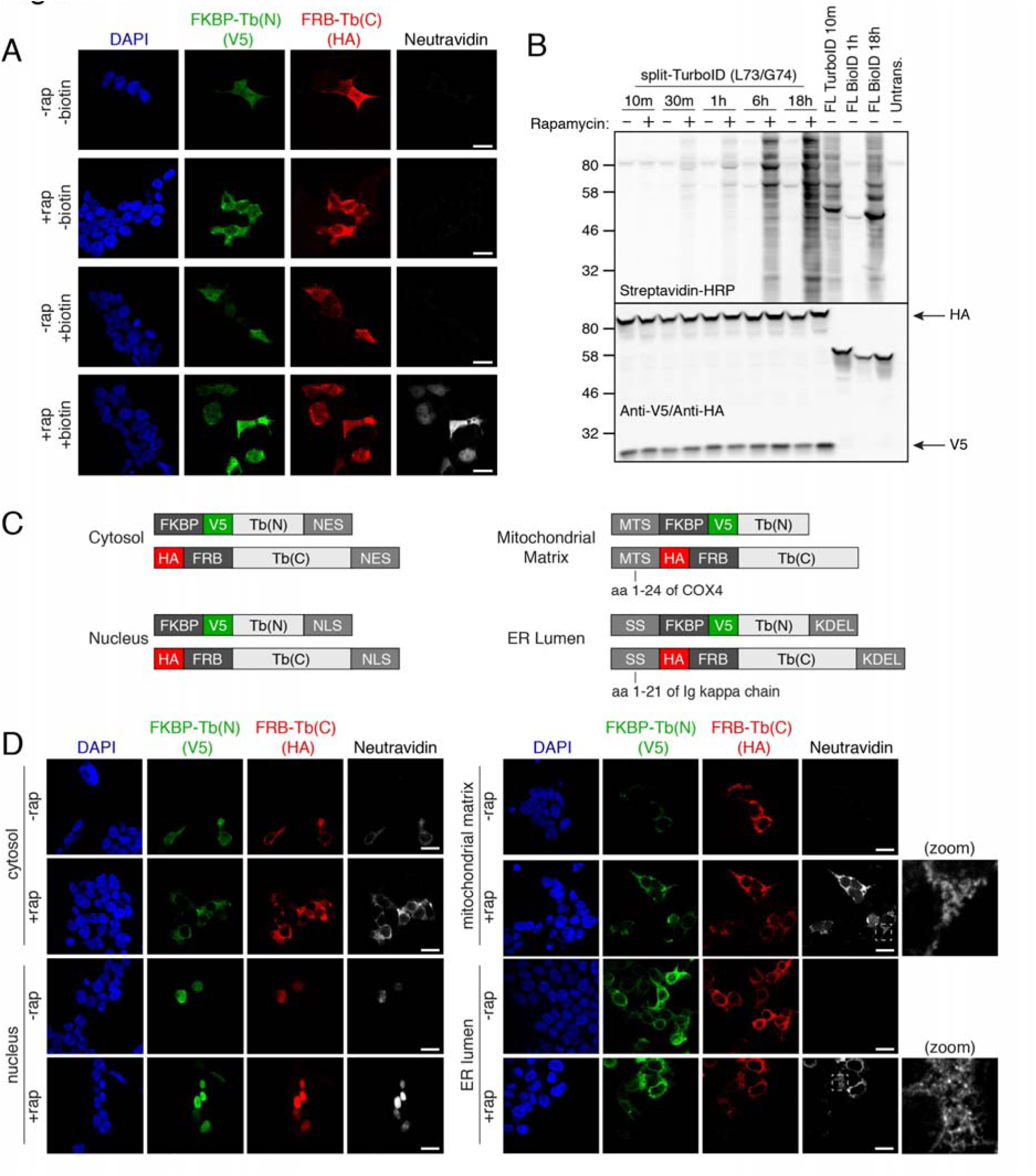
Characterization of low-affinity split-TurboID. *(A)* Confocal fluorescence imaging of low-affinity split-TurboID (split site L73/G74). HEK293T cells were transiently transfected and incubated with 50 μM biotin and 100 nM rapamycin for 1 hour, then fixed and stained with anti-V5 to detect the N-terminal fragment (Tb(N)), anti-HA to detect the C-terminal fragment (Tb(C)), and neutravidin-647 to detect biotinylated proteins. Scale bars, 20 μm. *(B)* Split-TurboID time course. HEK293T cells transiently transfected with split-TurboID constructs were treated with 50 μM biotin and 100 nM rapamycin for the indicated times, and whole cell lysates were analyzed by streptavidin blotting. Full-length TurboID (10 minutes) and full-length BioID (1 hour and 18 hours) were included for comparison. *(C)* Design of constructs used to target split-TurboID fragments to various cellular compartments. (MTS = mitochondrial targeting sequence, SS = signal sequence) *(D)* Confocal fluorescence imaging of low-affinity split-TurboID targeted to various cellular compartments. HEK293T cells were labeled and imaged as in (A). Fluorescence intensities are not normalized across cellular compartments. Zoomed images of the boxed regions are included. Scale bars, 20 μm.

We also generated constructs fusing split-TurboID with various localization sequences to target the fragments to different subcellular compartments (cytosol, nucleus, mitochondrial matrix, and ER lumen) (Figure 2C). Confocal fluorescence imaging of cells expressing these constructs labeled with rapamycin and biotin for 1 hour shows compartment-specific targeting and rapamycin-dependent biotinylation in all compartments tested (Figure 2D, Supplementary Figure 2B, C).

### Using split-TurboID for proximity labeling at ER-mitochondria contacts

ER-mitochondria contacts are important in a variety of biological processes, including Ca^+2^ signaling, lipid metabolism, nutrient signaling, and mitochondrial fission (15, 26, 27). There is tremendous interest in understanding the molecular composition of these contacts. Biochemical purification of mitochondria-associated membranes (MAMs) has frequently been used to study ER-mitochondria contacts (15), but MAMs encompass much more than just mitochondria-associated ER microsomes; they also include contaminants from the plasma membrane, Golgi, peroxisomes, and nuclear membrane (28). To provide a more specific alternative, we recently applied APEX proximity labeling to produce separate proteomic maps of the ER membrane and outer mitochondrial membrane, and then intersected the datasets to identify candidate ER-mitochondria contact residents (9). This resulted in the discovery of a novel ER-mitochondria tethering protein (SYNJ2BP), but many of the hits were merely dual-localized ER and mitochondria proteins.

We sought to use split-TurboID reconstitution across ER-mitochondria contacts in order to map this compartment directly, with much greater specificity than both MAM purifications and separate APEX tagging plus dataset intersection. To target split-TurboID to the ERM and OMM, we fused the fragments to the transmembrane domains of ERM-resident protein Cb5 and OMM-resident protein Tom20, respectively. We also included FKBP and FRB domains to enable rapamycin-induced heterodimerization (Figure 3A, B). In U2OS cells, we could observe some biotinylation activity in the absence of rapamycin, but it was substantially increased upon rapamycin addition (Figure 3C). Thus, it appears that close apposition of mitochondrial and ER membranes is sufficient to mediate some split-TurboID reconstitution, but rapamycin addition further enhances the reconstitution.

**Figure 3.**
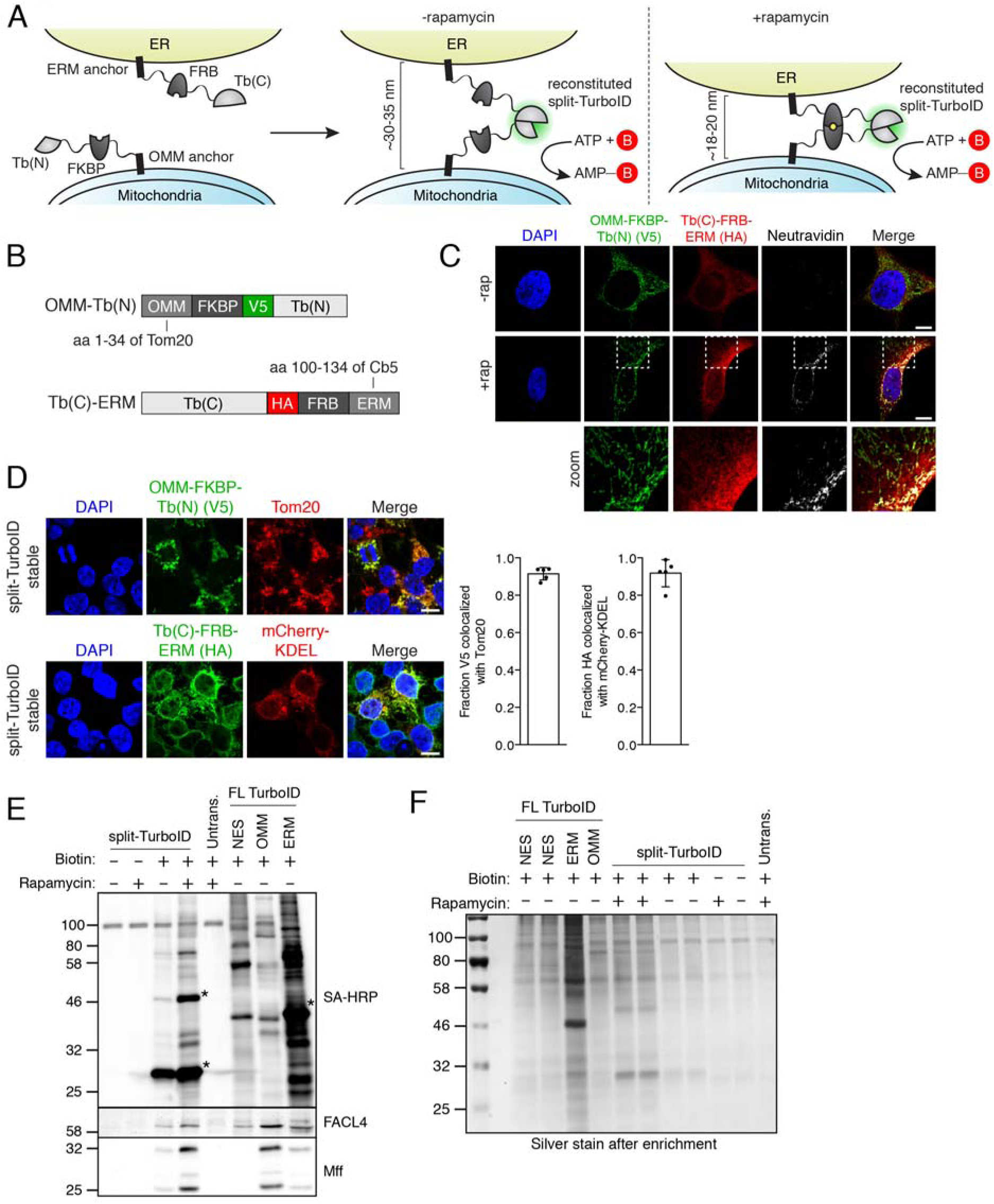
Reconstitution of split-TurboID at ER-mitochondria contact sites. *(A)* Schematic of split-TurboID reconstitution across ER-mitochondria contacts in the presence or absence of rapamycin for inducing dimerization. *(B)* Design of constructs targeting split-TurboID fragments to the outer mitochondrial membrane (OMM) and ER membrane (ERM). (OMM-targeted construct = 259 amino acids; ERM-targeted construct = 429 amino acids.) *(C)* Confocal fluorescence imaging of split-TurboID activity at ER-mitochondria contacts. Constructs were introduced into U2OS cells using lentivirus. Two days after transduction, cells were incubated with 50 μM biotin and 100 nM rapamycin for 1 hour, then fixed and stained with anti-V5 to detect the N-terminal fragment (Tb(N)), anti-HA to detect the C-terminal fragment (Tb(C)), and neutravidin-647 to detect biotinylated proteins. Zoomed images of the boxed regions are included. Scale bars, 20 μm. *(D)* Localization of split-TurboID in HEK293T cells stably expressing constructs from (B). Cells were fixed and stained with anti-V5 to detect the OMM-targeted N-terminal fragment (Tb(N)) or with anti-HA to detect the ERM-targeted C-terminal fragment (Tb(C)). Tom20 and mCherry-KDEL were used as mitochondrial and ER markers, respectively. Scale bars, 10 μm. Colocalization of V5 with Tom20 and HA with mCherry-KDEL are shown on the right. Quantitation from 5 fields of view per condition are included. *(E)* Enrichment of known ER-mitochondria proteins by split-TurboID-catalyzed proximity labeling. HEK293T cells stably expressing OMM/ERM-targeted split-TurboID constructs were treated with 50 μM biotin and 100 nM rapamycin for 4 hours. HEK293T cells stably expressing NES-, OMM-, or ERM-targeted full-length TurboID were treated with 50 μM biotin for 1 minute. Biotinylated proteins were enriched from lysates using streptavidin beads, eluted, and analyzed by streptavidin blotting. Streptavidin-enriched material was probed with antibodies against FACL4 and Mff. Asterisks indicate ligase self-biotinylation. *(F)* Enrichment of biotinylated proteins for proteomics. Samples were generated as in (E). Biotinylated proteins were enriched from lysates using streptavidin beads, eluted, and analyzed by silver stain.

### Split-TurboID mediated proteomic mapping of ER-mitochondria contacts

We designed our proteomic experiment to probe ER-mitochondria contact sites in HEK293T, both in the absence of rapamycin addition (when split-TurboID reconstitution is mediated only by native ERM and OMM proximity) and in the presence of rapamycin (which enhances split-TurboID reconstitution at ER-mitochondria contacts). We generated stable HEK293T cells expressing the split-TurboID constructs, or the reference constructs TurboID-NES (full-length TurboID in the cytosol), TurboID-OMM (full-length TurboID on the outer mitochondrial membrane facing cytosol), and ERM-TurboID (full-length TurboID on the ER membrane facing cytosol). In HEK293T cells stably expressing split-TurboID or full-length TurboID, constructs displayed correct localization to mitochondria and ER organelles (Figure 3D, Supplementary Figure 4A-C). For split-TurboID, biotinylation activity was again observed in the absence of rapamycin, but was substantially increased upon rapamycin addition (Supplementary Figure 3A).

Due to the difference in activity levels, the split-TurboID samples were treated with biotin (with or without rapamycin) for 4 hours, while the full-length TurboID samples were labeled for only 1 minute. Under these conditions, we observed comparable levels of biotinylated proteins in our split-TurboID and full-length TurboID samples both before and after streptavidin bead enrichment (Figure 3F, Supplementary Figure 3D, E). We also verified that these labelling conditions did not perturb organelle morphology or artificially increase ER-mitochondria contacts in our stable cells (Supplementary Figure 3B, C). To test if OMM/ERM-targeted split-TurboID could preferentially enrich known ER-mitochondria contact site proteins, we performed Western blot analysis of the streptavidin-enriched material. Figure 3E shows enrichment of the known ER-mitochondria contact proteins FACL4 and Mff (15, 17, 29) in split-TurboID samples, but not in TurboID-NES samples.

Next, we performed mass spectrometry on our streptavidin-enriched samples. After on-bead digestion of the protein samples, we labeled the peptides with isotopically distinct TMT (tandem mass tag) labels, enabling us to quantify the relative abundance of each protein across samples (Figure 4A). Input levels per sample were normalized prior to analysis, and we found that replicate samples were highly correlated (Supplementary Figure 5A-D). We analyzed our ERM and OMM datasets obtained from full-length TurboID using a ratiometric approach as previously described (30) and found our datasets to be highly specific both when compared to previously published datasets (7, 9) and when performing GO term enrichment analysis (Supplementary Figure 5E-P, Supplementary Figure 6A-E).

**Figure 4.**
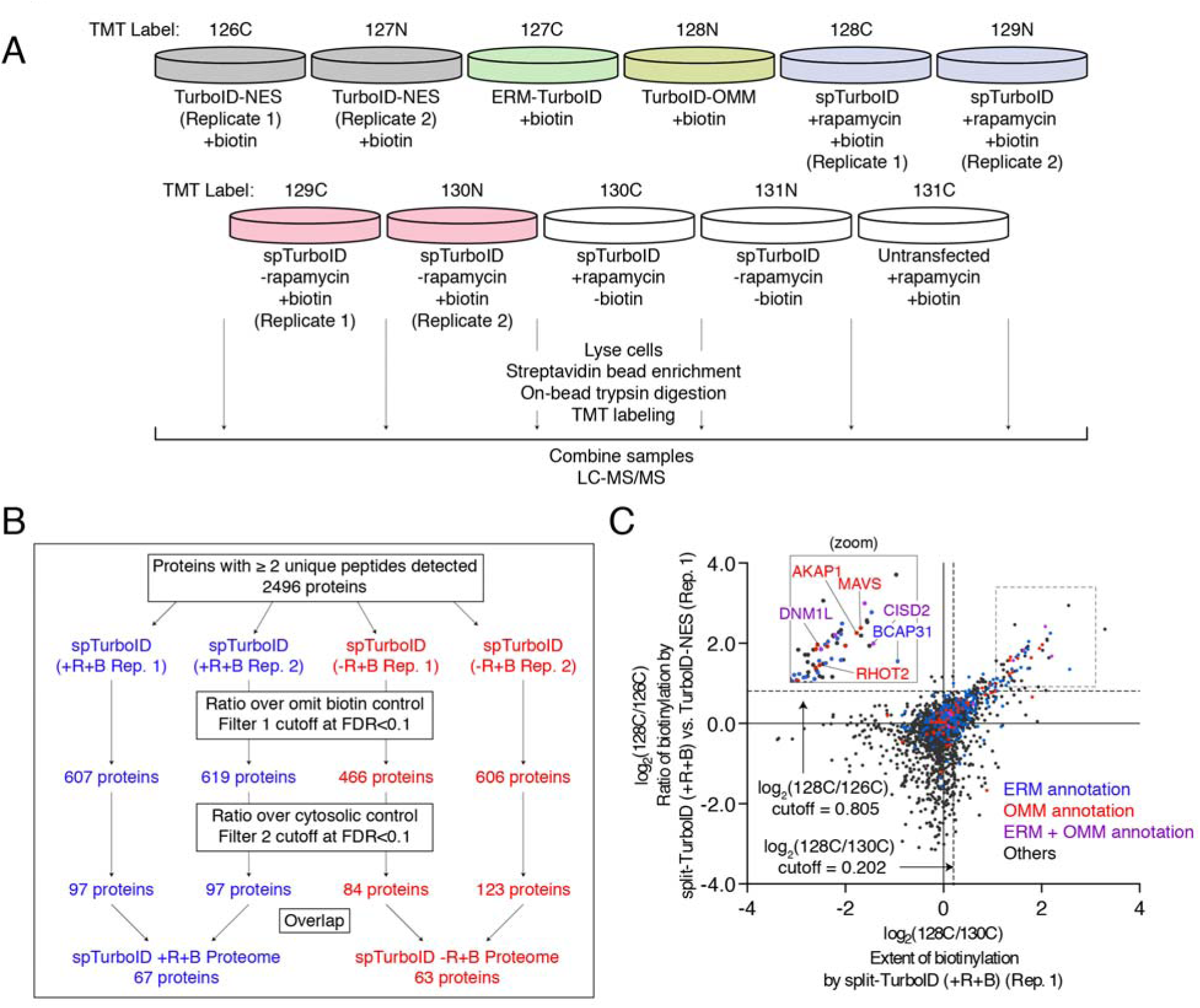
Proteomic mapping of ER-mitochondria contacts in HEK293T cells. *(A)* Experimental design and labeling conditions for MS-based proteomics. Cells stably expressing the indicated constructs were labeled with 50 μM biotin and 100 nM rapamycin. Split-TurboID (ERM/OMM) samples were labeled for 4 hours and full-length TurboID samples were labeled for 1 minute. Cells were then lysed, and biotinylated proteins were enriched using streptavidin beads, digested to peptides, and conjugated to TMT (tandem mass tag) labels. All samples were then combined and analyzed by LC-MS/MS. *(B)* Filtering scheme for mass spectrometric data. +R and -R refer to rapamycin, and +B and -B refer to biotin. For each dataset, proteins were first ranked by the extent of biotinylation (ratiometric analysis referencing omit biotin controls, filter 1). Next, proteins were ranked by relative proximity to ER-mitochondria contacts versus cytosol (ratiometric analysis referencing TurboID-NES, filter 2). *(C)* Scatterplot showing log_2_(128C/126C) (filter 1) versus log_2_(128C/130C) (filter 2) for each protein in replicate 1 of split-TurboID treated with rapamycin and biotin. Known ERM and OMM proteins (as annotated by GOCC) are labeled blue and red, respectively; ERM and OMM dual-annotated proteins are colored purple, and all other proteins are shown in black. Cutoffs used to filter the mass spectrometric data and obtain the filtered proteome are shown by dashed lines. Zoom of boxed region shown in the upper left.

For the analysis of our split-TurboID proteomic datasets, we began with 2496 detected proteins with two or greater unique peptides. We then took the replicates of each experimental condition (+rapamycin or -rapamycin) and applied two sequential filtering steps. First, the data was filtered by the extent of biotinylation by split-TurboID, where we established a cutoff at a 10% false discovery rate (FDR) for detection of mitochondrial matrix false positive proteins, referencing the “omit biotin” negative control samples. Second, the data were filtered further by the extent of biotinylation by split-TurboID at ER-mitochondria contacts relative to proteins biotinylated by TurboID in the cytosol, where we established a cutoff at a 10% FDR for detection of cytosolic false positive proteins (Figure 4B, Supplementary 7A-H). Applying both filters enriched for proteins with prior ERM and OMM annotation, as well as proteins that have been previously associated with ER-mitochondria contacts, including DNM1L, BCAP31, MAVS, and AKAP1 (16, 29, 31, 32) (Figure 4C). After filtering, we reached proteome lists of 67 proteins (+rapamycin list) and 63 proteins (-rapamycin list), 29 proteins of which were found in both lists (Figure 4B, Figure 5A).

**Figure 5.**
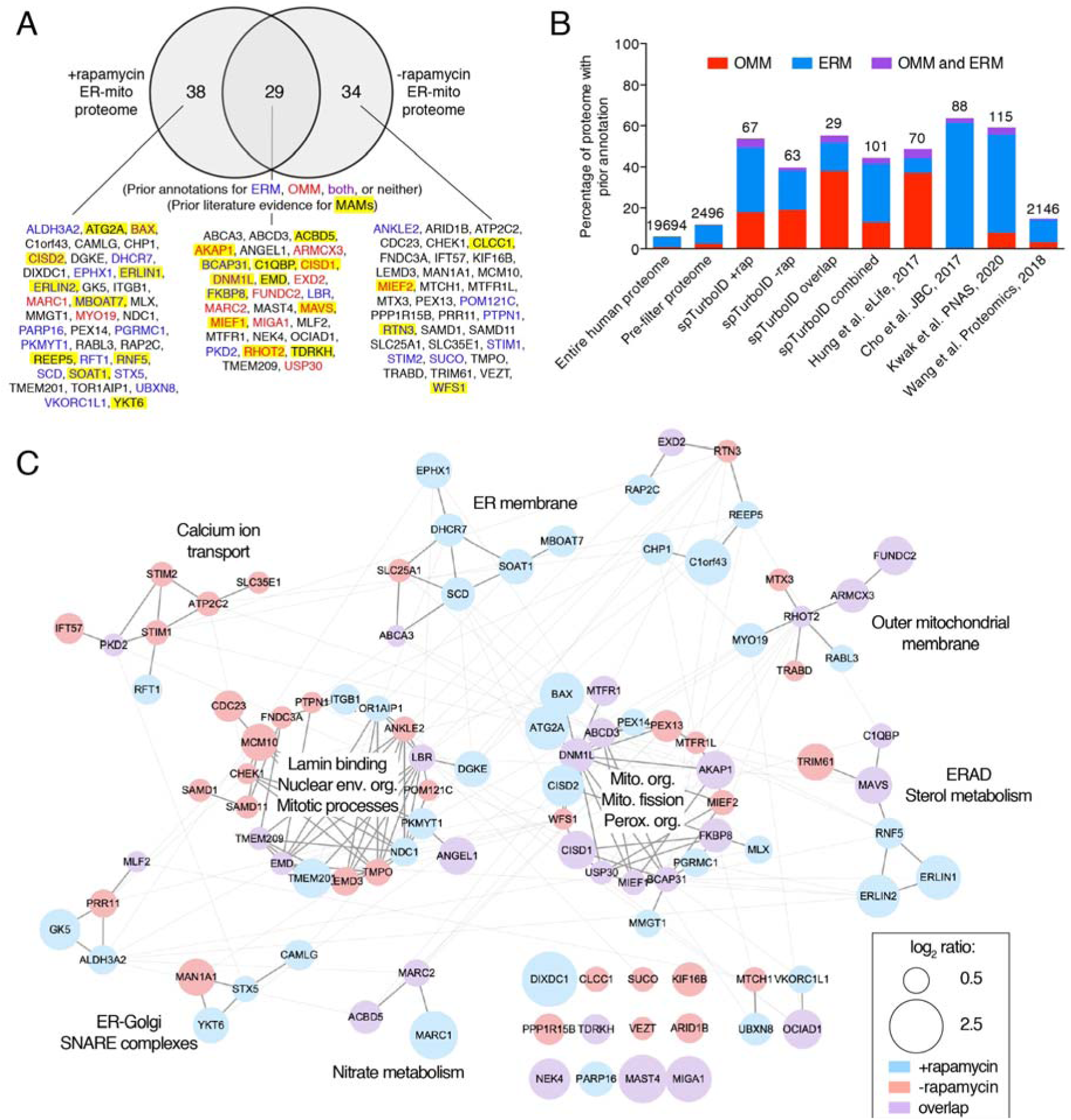
Analysis of ER-mitochondria contact proteomic dataset. *(A)* Venn diagram comparing proteome lists using split-TurboID +rapamycin versus-rapamycin. Proteins that were previously annotated ERM, OMM, both, or neither were labeled blue, red, purple, or black, respectively. Proteins with previous literature evidence for association with MAMs are highlighted in yellow. *(B)* Specificity analysis for proteomic datasets generated using split-TurboID compared to the entire human proteome and to previous datasets studying ER-mitochondria contacts or MAMs. Bar graph shows the percentage of each proteome with prior OMM annotation only, ERM annotation only, or both OMM and ERM annotation. Each bar is labeled with the size of the proteome. Hung et al. used APEX (9), Cho et al. used APEX and MAM fractionation (36), Kwak et al. used Contact-ID (21), and Wang et al. used MAM purification (35). *(C)* Markov clustering of split-TurboID proteome using protein-protein interaction scores from the STRING database (33). Gray lines denote protein-protein interactions. Nodes are colored based on whether the corresponding protein was found in the +rapamycin proteome, -rapamycin proteome, or both proteomes. Node size is correlated with the relative enrichment of each protein at ER-mitochondria contacts versus cytosol (log_2_(128C/126C) or log_2_(129C/126C)). Each cluster is labeled with associated GO terms.

GO term enrichment analysis of each list largely recovered ER and mitochondria membrane-associated terms, suggesting high specificity (Supplementary Figure 9A, B). We calculated the fraction of proteins in each dataset with prior ERM or OMM annotation, and arrived at 44% for our combined dataset (proteins present in either +rapamycin or -rapamycin lists), and 55% for our overlapping dataset (proteins present in both lists), both much greater than the equivalent percentages for the entire human proteome (6%) or our pre-filter proteome (11%). We next performed Markov clustering of our proteomic datasets, using known protein-protein interactions from the STRING database (33). Figure 5C shows the resulting network and GO terms associated with each cluster. The GO terms mitochondrial organization, mitochondrial fission, sterol metabolism, and calcium ion transport are consistent with known roles of ER-mitochondria contacts (15, 27, 34).

We compared our split-TurboID proteomes to previous ER-mitochondria contact proteomes obtained by other methods. The four comparison datasets were: (1) a MAM preparation from human tissue (35), (2) a study combining MAM isolation and proximity labeling (36), (3) our previous study using APEX labeling on the OMM and ERM separately, followed by dataset intersection (9), and (4) the Contact-ID-generated ER-mitochondria proteome (21). Figure 5B shows that specificity, as measured by the fraction of proteins with prior OMM or ERM annotation, is similar for all of the proximity labeling-based studies, but much poorer for the MAM dataset, which contains contaminants from many other organelles. To quantify sensitivity, we compiled a list of 20 human proteins with prior literature evidence of ER-mitochondria contact localization. We detected only 2 of these proteins, similar to other proximity labeling studies, while the MAM dataset recovered much more. Overall, MAM purifications are more sensitive, but at the expense of specificity. Proximity labeling has the opposite characteristics – high specificity, but poorer recovery, particularly for dual-localized proteins which are known to be removed by the ratiometric filtering process (for example, a protein dual-localized to ER-mitochondria contacts *and* the cytosol would be removed in the second ratiometric filtering step using cytosolic TurboID-NES as a reference).

### Validation of ER-mitochondria contact “orphans”

Of the 29 proteins detected in both +rapamycin and -rapamycin datasets, 12 have previously been detected in MAMs or localized to ER-mitochondria contacts in literature. The remainder are “orphans”, or proteins with no prior literature connection to ER-mitochondria contacts. To determine if these are bona fide ER-mitochondria contact residents or false positives, we selected 6 proteins for which high quality commercial antibodies exist, and blotted for their presence in purified MAMs. All 6 were found to be enriched in MAMs as well as in other compartments consistent with their literature annotation (e.g. MAM + ER for LBR; MAM + mitochondria for EXD2) (Figure 6A).

**Figure 6.**
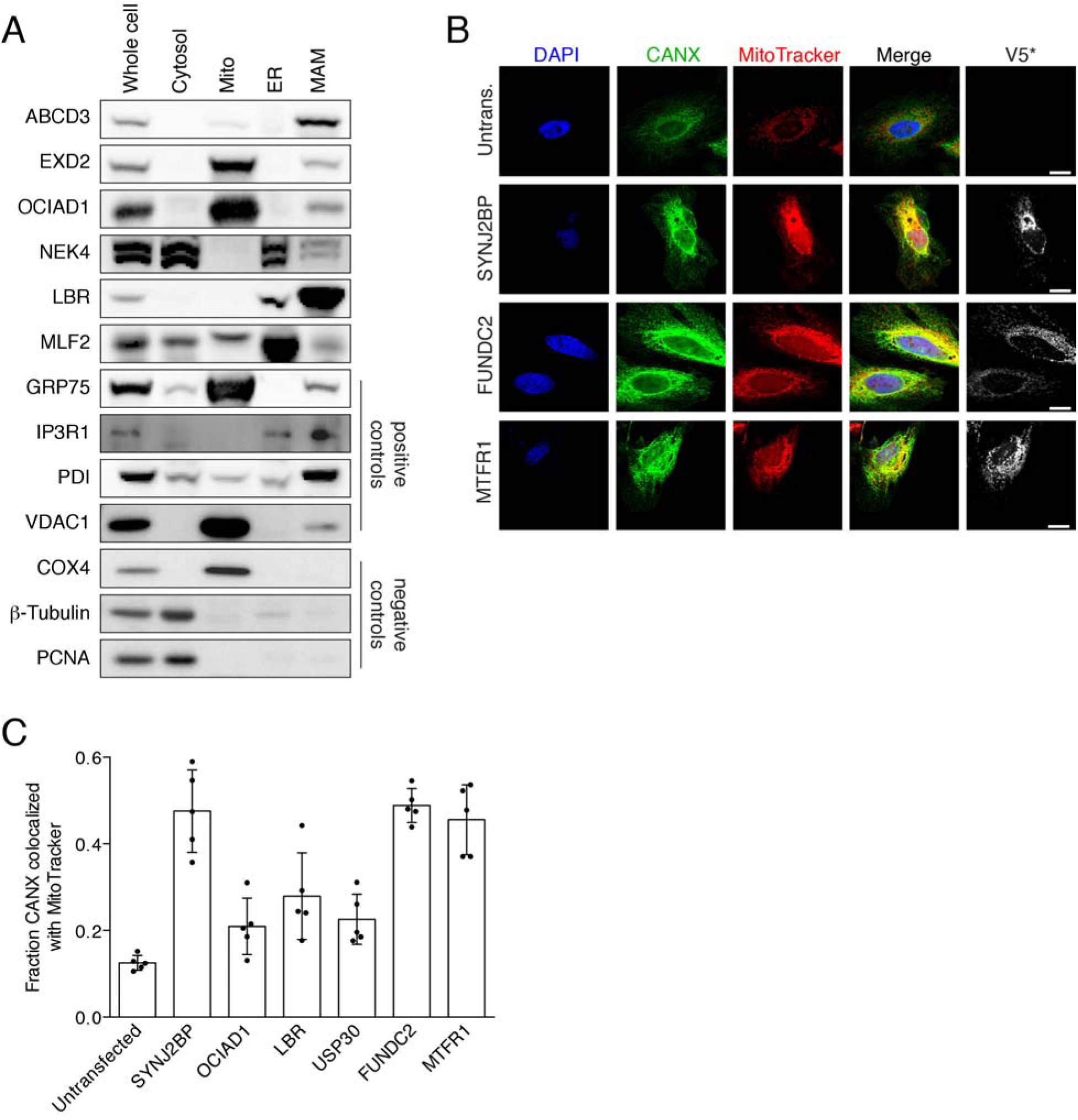
Validation of proteomic hits in MAMs and by imaging. *(A)* Western blotting of candidate ER-mitochondria contact proteins ABCD3, EXD2, OCIAD1, NEK4, LBR, and MLF2 in MAM fractions. HEK293T cells were collected and subjected to subcellular fractionation to obtain cytosol, mitochondria (mito), endoplasmic reticulum (ER), and mitochondria-associated membrane (MAM) fractions. GRP75, IP3R1, PDI, and VDAC1 are known MAM proteins and were included as positive controls. Negative controls are COX4 (inner mitochondrial membrane), β-tubulin (cytosol), and PCNA (nucleus). *(B)* Overexpression imaging assay. Candidate ER-mitochondria contact proteins FUNDC2 and MTFR1(each V5-tagged) were overexpressed in HeLa cells. V5-tagged SYNJ2BP (9) was included as a positive control. MitoTracker stains mitochondria and anti-CANX antibody stains ER. *(C)* Quantification ER-mitochondria overlap (colocalization of CANX and MitoTracker signals) in (B) and Supplementary Figure 9F. Five fields of view were analyzed per condition.

In addition to MAM blotting, we selected a subset of orphans for functional analysis. Previously, we showed that overexpression of the ER-mitochondria tether SYNJ2BP in HeLa cells causes a dramatic increase in the extent of overlap between ER and mitochondria by imaging (9). We V5-tagged the proteins FUNDC2, LBR, MTFR1, OCIAD1, and USP30 and overexpressed them in HeLa cells, which were then stained for mitochondria and ER markers. Confocal imaging shows that FUNDC2 and MTFR1, along with the positive control SYNJ2BP, causes a significant increase in colocalization of ER and mitochondria markers, compared to untransfected control HeLa cells (Figure 6B, C). The other three proteins tested did not give this phenotype (Supplementary Figure 9F). FUNDC2 and MTFR1 may have tethering functions at ER-mitochondria contacts that are upregulated in this gain-of-function assay.

### Cell-cell contact dependent reconstitution of split-TurboID

In addition to permitting PL with greater spatial specificity, split-TurboID could potentially enable conditional PL dependent upon specific inputs or signaling events. For instance, in neuroscience, immunology, and cancer biology, there is great interest in characterizing the transcriptomes and proteomes of cell subpopulations that have made contact with specific “sender” cells (for example, neurons downstream of specific pre-synaptic inputs, or immune cells that come into contact with a tumor cell). If we could drive intracellular split-TurboID reconstitution, specifically in “receiver” cell populations that come into contact with defined “sender” cells, then this could potentially enable PL-based proteomic analysis of functionally relevant cellular subpopulations.

To test this, we designed a signaling network that utilizes the trans-cellular interaction between glucagon peptide and its receptor (GCGR), employed in the trans-synaptic tracing tool trans-Tango (37). In our design, “receiver” cells express glucagon receptor (GCGR) and arrestin, each fused to a split-TurboID fragment. Upon cell-cell contact with “sender” cells expressing surface glucagon peptide, GCGR is activated, resulting in arrestin recruitment and reconstitution of split-TurboID (Figure 7A). Confocal fluorescence imaging of receiver and sender cell co-cultures after biotin treatment for 1 hour showed highly specific localization of biotinylated proteins, specifically at sender:receiver cell-cell contacts (red:green overlap sites in Figure 7B).

**Figure 7.**
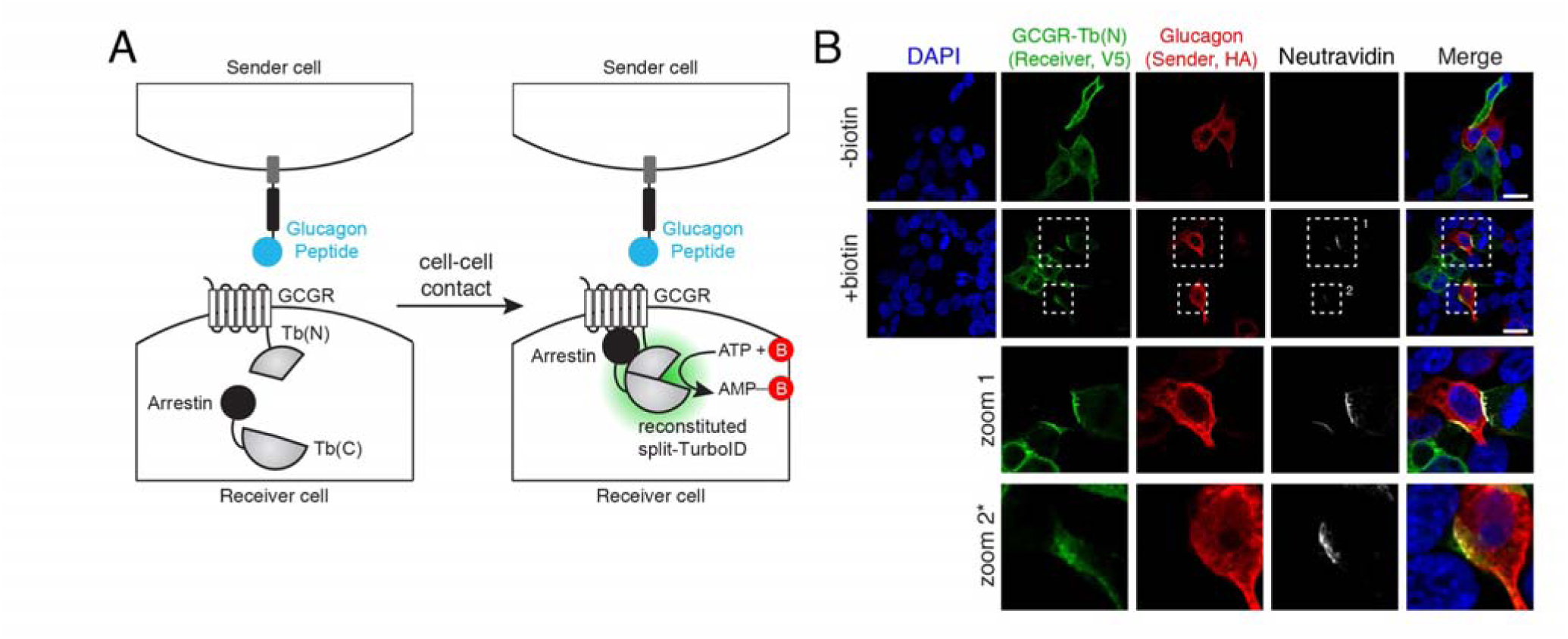
Cell-cell contact dependent reconstitution of split-TurboID. *(A)* Schematic of cell-cell contact dependent split-TurboID reconstitution. *(B)* Confocal fluorescence imaging of “sender” cells expressing cell surface targeted glucagon peptide co-cultured with “receiver” cells expressing split-TurboID fragments. Scale bars, 20 μm. The contrast has been increased in the second zoom box.

## Discussion

We have engineered both high-affinity and low-affinity forms of split-TurboID, which are simpler to use and much more active than previously reported split enzymes for PL (18–21). We show that reconstitution is fast and effective in multiple cellular subcompartments (Figure 2D). Low-affinity split-TurboID reconstitution can be driven by a drug, protein-protein interaction, organelle-organelle contact, or cell-cell contact.

We used split-TurboID for proteomic mapping of ER-mitochondria contact sites and identified 67 proteins in our +rapamycin dataset and 63 proteins in our-rapamycin dataset which were reproducibly enriched over controls. While these lists contain many proteins that were previously detected in MAMs, there are more than 70 “orphans” with no prior literature connection to ER-mitochondria contacts. One interesting example is MIGA1, or mitoguardin 1, identified in both our +rapamycin and -rapamycin datasets. MIGA1 has been previously shown to function downstream of the mitofusins and regulate mitochondrial fusion (38). A prior study showed that MIGA1 localizes to the OMM but not specifically to ER-mitochondria contact sites (38). Interestingly, MIGA2, which is highly related to MIGA1, was recently shown to bind VAPA and VAPB and link ER-mitochondria contacts to lipid droplets to promote adipocyte differentiation (39).

We validated eight split-TurboID-identified ER-mitochondria contact orphans, using biochemical fractionation and overexpression imaging. These proteins span a range of functions and their assignment to ER-mitochondria contacts has interesting biological implications. For instance, ABCD3 is a known peroxisomal membrane protein that has been shown to be a transporter of branched chain fatty acids into the peroxisome (40, 41). Our finding that ABCD3 localizes to ER-mitochondria contacts is consistent with previous studies suggesting that peroxisome biogenesis may occur at ER-mitochondria contacts (42, 43). EXD2 is an endonuclease that has been shown to localize to both mitochondria and the nucleus (44, 45); its localization to ER-mitochondria contacts establishes a possible link between mitochondrial function and nuclear DNA maintenance, which has also been suggested in previous work (45). LBR is a lamin B receptor that has been localized to the nuclear envelope and is involved in nuclear envelope disassembly during mitosis (46). Our finding that LBR localizes to ER-mitochondria contacts (Figure 6A), along with our analysis showing that a number of proteins involved in lamin binding, nuclear envelope organization, and mitosis may be enriched at ER-mitochondria contact sites (Figure 5C), suggest that these processes may be regulated in part by

ER-mitochondria contact. MTFR1 and FUNDC1 are the two split-TurboID-identified orphans that caused an increase in ER-mitochondria overlap upon overexpression (Figure 6B, C). While MTFR1 has been shown to regulate mitochondrial dynamics (47), and FUNDC1, related to FUNDC2, has been shown to mediate hypoxia-mediated mitophagy (48), MTFR1 and FUNDC2 have not previously been localized to ER-mitochondria contacts or shown to have organelle tethering activity.

Comparing our +rapamycin and -rapamycin datasets, we observed somewhat more proteins with prior ER, mitochondria, and MAM annotations in the +rapamycin condition. Because in our construct design, FRB and FKBP also function like linkers that extend the reach of the split-TurboID fragments, we hypothesize that the +rapamycin proteome may be probing closer ER-mitochondria contacts, while the -rapamycin proteome may also be probing wider contacts by reaching farther as shown in Figure 3A. Perhaps MAM preparations in previous studies are biased towards closer ER-mitochondria contacts if they are more likely to survive the fractionation process while wider contacts may be under-represented. Thus, while there is less prior annotation information for proteins in the -rapamycin proteome, these proteins may still be bona fide ER-mitochondria contact proteins.

In addition to generating ER-mitochondria contact candidates, our proteomic experiment also produced OMM and ERM proteomes, via the full-length TurboID constructs we targeted to OMM and ERM, respectively. Using these datasets, we could categorize our ER-mitochondria contact hits as additionally localized to OMM and/or ERM, or not (Supplementary Figure 9D; each protein is classified as being a resident of “Contacts only”, “Contacts and OMM”, “Contacts and ERM”, or “Contacts, OMM, and ERM”). The categorizations based on our proteomic data are largely consistent with prior literature, and with our fractionation blotting data (Figure 6A-C, Supplementary Figure 9E, F). For several proteins, such as OCIAD1 and NEK4, our proteomic data newly assign them to mitochondrial and ER compartments of the cell.

When comparing our proteome with that obtained via Contact-ID (21), we find that both proteomes detected a similar number of proteins (101 proteins using split-TurboID and 115 proteins using Contact-ID). Of the eight proteins we validated, four (FUNDC2, EXD2, OCIAD1, and LBR) were also detected using Contact-ID (Supplementary Figure 8A-C). Conversely, FKBP8, which the authors of Contact-ID show facilitates ER-mitochondria contact formation and local calcium transport (21), was also present in our proteome. While both proteomes have high specificity, measured by the fraction of proteins with prior OMM or ERM annotation, we find that the proteome generated from Contact-ID is more biased towards ER membrane proteins, whereas we observe more balance between OMM and ERM proteins in our split-TurboID proteome (Figure 5B). This difference may possibly arise from the longer labeling period used for Contact-ID (16 hours vs. 4 hours for split-TurboID).

Overall, we have developed a split-TurboID tool that can be conditionally reconstituted for spatially-specific proximity labeling in cells. Especially when combined with functional assays and screens, split-TurboID-based PL can be a powerful tool for biological discovery around organelle contact sites or macromolecular complexes. Split-TurboID may also improve signal-to-noise for challenging targeting applications, such as dCas9-directed PL of specific genomic loci (49, 50), or dCas13-directed PL of specific cellular RNAs (51).

## Acknowledgements

This work was supported by the NIH R01-DK121409 (to A.Y.T. and S.A.C.), Stanford Wu Tsai Neurosciences Institute Big Ideas (to A.Y.T.), Korea Health Technology R&D Project Grant KHIDI HI16C1501 (to I.K.L.), National Research Foundation of Korea (NRF) grants 2017R1A2B3006406 (to I.K.L.) and 2019R1A2C3008463 (to H.W.R.), and Organelle Network Research Center grant NRF-2017R1A5A1015366 (to H.W.R.). K.F.C. was supported by NIH Training Grant 2T32CA009302-41 and the Blavatnik Graduate Fellowship. T.C.B. was supported by Dow Graduate Research and Lester Wolfe Fellowships. A.Y.T. is an investigator of the Chan Zuckerberg Biohub.

## Author Contributions

K.F.C. and A.Y.T. designed the research and analyzed all the data unless otherwise stated. K.F.C. performed all experiments unless otherwise stated. K.F.C. and T.C.B. designed split-TurboID constructs for screening. K.F.C. and S.R. performed molecular cloning for testing split-TurboID. K.F.C., T.C.B., A.Y.T., N.D.U., T.S., and S.A.C. designed the proteomics experiments. K.F.C. prepared the proteomic samples. N.D.U. and T.S. processed the proteomic samples and performed mass spectrometry. T.T. and I.K.L. performed subcellular fractionation. T.T. and K.F.C performed Western blotting for MAM proteins. K.F.C. and A.Y.T. wrote the manuscript with input from all other authors.

## Methods

### Cloning

All constructs were generated using standard cloning techniques. PCR fragments were amplified using Q5 polymerase (NEB). Vectors were digested using enzymatic restriction digest and ligated to gel purified PCR products using Gibson assembly. Ligated plasmid products were transformed into competent XL1-Blue bacteria.

### Split site pairs selection

For selecting split site pairs for testing for split-TurboID, split sites tested in the study developing split-BioID (20) were mapped onto TurboID: 256/257, 258/259, 270/271, 273/274. We also applied the SPELL algorithm (23) to wild-type *E. coli* biotin ligase (BirA; PDB: 1HXD (24) and PDB: 2EWN (25)), from which TurboID is derived. We selected 5 split site pairs predicted by SPELL for testing. Due to a numbering error during cloning, the 5 split site pairs tested were 73/74, 98/99, 101/102, 112/113, 212/213. To further optimize our split-TurboID tool, we also selected split site pairs near our candidate split site: 71/72, 72/73, 74/74, 75/76.

### Mammalian cell culture, transfection, and stable line generation

HEK293T cells from ATCC were cultured as a monolayer in Dulbecco’s Modified Eagle’s Medium (DMEM) with 4.5 g/L glucose and L-glutamine supplemented with 10% (w/v) fetal bovine serum, 50 units/mL penicillin, and 50 μg/mL streptomycin at 37°C under 5% CO_2_. For confocal imaging experiments, glass coverslips were coated with 50 μg/mL fibronectin in DPBS for at least 20 minutes at room temperature before plating; cells were grown on glass coverslips in 24-well plates with 500 μL growth medium. For Western blot experiments, cells were grown in 6-well plates with 2 mL growth medium.

For transient expression, cells were transfected at approximately 50% confluency using 5 μL/mL Lipofectamine2000 and 250 ng/mL plasmid in serum-free media. To generate lentiviruses, HEK293T cells were cultured in T25 flasks and transfected at approximately 60% confluency with 2500 ng of the lentiviral vector containing the gene of interest and lentiviral packaging plasmids pVSVG (250 ng) and Δ8.9 (2250 ng) with 30 μL polyethyleneimine (PEI) in water (pH 7.3, 1 mg/mL) in serum-free media. After 48 hours, the cell medium containing the lentivirus was harvested and filtered using a 0.45 μm filter.

For generation of stable cell lines, HEK293T cells were infected with lentivirus at approximately 50%, followed by selection with 8 μg/mL blasticidin (at least 4 days) and 250 μg/mL hygromycin (at least 7 days) in growth medium before use in experiments.

### Biotin labeling with TurboID and split-TurboID

For biotin labeling of transiently transfected cells, biotin and rapamycin were added 18 hours following transfection. For full-length TurboID, cells were treated with 50 μM biotin and incubated at 37°C; for split-TurboID, cells were treated with 50 μM biotin and 100 nM rapamycin and incubated at 37°C. Labeling was stopped after the indicated time periods by transferring cells to ice and washing with cold PBS.

### Protein gels and Western blots

HEK293T cells expressing indicated constructs were plated, transfected, and labeled as previously described. After labeling, cells were transferred to ice and washed twice with cold PBS. Cells were then detached from the well or flask by pipetting a stream of cold PBS directly onto cells. The cells were the collected and pelleted by centrifugation at 2,500 rpm for 3 minutes at 4°C. The supernatant was removed, and the pellet was lysed by resuspending in RIPA lysis buffer (50 mM Tris pH 8, 150 mM NaCl, 0.1% SDS, 0.5% sodium deoxycholate, 1% Triton X-100, 1x protease inhibitor cocktail (Sigma-Aldrich), and 1 mM PMSF) and incubating for 10 minutes at 4°C. Lysates were clarified by centrifugation at 12,000 rpm for 10 minutes at 4°C. Samples were mixed with loading buffer prior to loading and separation on 9% SDS-PAGE gels. Silver-stained gels were generated using the Pierce Silver Stain kit (ThermoFisher). For Western blotting, proteins separated on SDS-page gels were transferred to nitrocellulose membranes and stained with Ponceau S (2 minutes in 0.1% (w/v) Ponceau S in 5% acetic acid/water). The blots were then blocked in 5% milk (w/v) in TBST (Tris-buffered saline, 0.1% Tween 20) for 1 hour at room temperature. Blots were washed twice with TBST then incubated with primary antibodies in 3% BSA (w/v) in TBST overnight at 4°C. Blots were then washed three times with TBST for 5 minutes each, and incubated with secondary antibodies or 0.3 μg/mL streptavidin-HRP in 3% BSA (w/v) in TBST for 1 hour at 4°C. The blots were washed four times with TBST for 5 minutes each before developing with Clarity Western ECL Blotting Substrates (Bio-Rad) and imaging. Blots were imaged on a ChemiDoc XRS+ imager (Bio-Rad) or an Odyssey CLx imager (LI-COR). Quantitation of Western blots was performed using ImageJ on raw images.

### Cell culture fixation, staining, and confocal imaging

HEK293T cells expressing indicated constructs were plated onto coverslips, transfected, and labeled as previously described. After labeling, cells were fixed with 4% (v/v) paraformaldehyde diluted in serum-free media and 20% (v/v) 5x PHEM buffer (300 mM PIPES, 125 mM HEPES, 50 mM EGTA, 10 mM MgCl_2_, 0.6 M sucrose, pH 7.3) for 10 minutes. The solution was removed and cells were then permeabilized with cold methanol at 4°C for 10 minutes. Cells were then washed three times with PBS and blocked in 1% BSA (w/v) in PBS for 30 minutes at room temperature. Cells were incubated with primary antibody in 1% BSA (w/v) in PBS for 3 hours at room temperature. After washing three times with PBS, cells were incubated with DAPI, secondary antibodies, and neutravidin-Alex Fluor 647 in 1% BSA (w/v) in PBS for 1 hour at room temperature. Cells were then washed with PBS three times, mounted onto glass slides, and imaged by confocal fluorescence microscopy. Confocal imaging was performed with a Zeiss AxioObserver inverted microscope with a 63x oil-immersion objective. The following combinations of laser excitation and emission filters were used for various fluorophores: DAPI (405 nm laser excitation, 445/40 nm emission), AlexaFluor 488 (491 nm laser excitation, 528/38 nm emission), AlexaFluor 568 (561 nm laser excitation, 617/73 nm emission), and AlexaFluor 647 (647 nm laser excitation, 680/30 nm emission). All images were collected with SlideBook (Intelligent Imaging Innovations) and processed with ImageJ.

For colocalization analysis, masks were generated in the channels of interest using a threshold value of >10x the background signal, unless otherwise noted, determined by measuring the intensity of a region outside a cell. Colocalization was quantified by measuring the number of pixels that surpassed the threshold in both channels. For colocalization of HA and mCherry-KDEL in Figure 3D, 3 mCherry-KDEL positive cells were analyzed per FOV.

### Sample preparation for proteomics

For each sample, HEK293T cells were cultured as previously described in T150 flasks. All cells used in proteomics experiments were stably expressing the indicated constructs. Split-TurboID samples were labeled with 50 μM biotin and 100 nM rapamycin for 4 hours, and full-length TurboID samples were labeled with 50 μM biotin for 1 minute. Labeling was stopped by placing cells on ice and washing with cold PBS twice. Cells were then detached from the well or flask by pipetting a stream of cold PBS directly onto cells. The cells were the collected and pelleted by centrifugation at 2,500 rpm for 3 minutes at 4°C. The supernatant was removed, and the pellet was lysed by resuspending in RIPA lysis buffer and incubating for 10 minutes at 4°C. Lysates were clarified by centrifugation at 12,000 rpm for 10 minutes at 4°C.

For enrichment of biotinylated proteins, 300 μL streptavidin-coated magnetic beads (Pierce) were washed twice with RIPA lysis buffer and incubated with clarified lysates for each sample with rotation at 4°C overnight. The beads were then washed twice with 1 mL of RIPA lysis buffer, once with 1 mL 1M KCl, once with 1 mL 0.1M Na_2_CO_3_, once with 1 mL 2M urea in 10 mM Tris-HCl (pH 8.0), and twice with 1 mL RIPA lysis buffer. The beads were subsequently washed with 1 mL digestion buffer (75 mM NaCl, 1 mM EDTA, 50 mM Tris-HCl, pH 8.0) twice. The beads were then resuspended in 80 μL digestion buffer, transferred to a new Eppendorf tube, frozen on dry ice, and delivered to Steve Carr’s laboratory (Broad Institute) for further processing and preparation for LC-MS/MS analysis.

For each proteomic sample, 0.2% of the lysate was removed prior to enrichment to verify construct expression and confirm successful biotinylation as shown in Figure 4D. After enrichment, 5% of beads were removed and biotinylated proteins were eluted by boiling the beads in 80 μL 3x protein loading buffer supplemented with 20 mM DTT and 2 mM biotin. The eluted proteins were separated by SDS-PAGE gel and analyzed by Western blotting (1.25%) and Silver stain (3.75%) to verify successful enrichment of biotinylated proteins (Figure 4D).

### On-bead trypsin digestion of biotinylated proteins

For sample preparation, proteins bound to streptavidin beads were washed twice with 200 μL 50 mM Tris-HCl (pH 7.5) and then twice with 2M urea in 50 mM Tris-HCl (pH 7.5). The buffer was removed and beads were incubated with 80 μL 2M urea in 50 mM Tris-HCl (pH 7.5) containing 1 mM DTT and 0.4 μg trypsin for 1 hour at 25°C with shaking. After 1 hour, the supernatant was removed and transferred to a fresh tube. The remaining streptavidin beads were washed twice with 60 μL 2M urea in 50 mM Tris-HCl (pH 7.5) and the washes were combined with the previous on-bead digest supernatant. The resulting mixture was reduced with 4 mM DTT for 30 minutes at 25°C with shaking. The samples were then alkylated with 10 mM iodoacetamide for 45 minutes in the dark at 25°C with shaking. An additional 0.5 μg trypsin was added to each sample and incubated overnight at 25°C with shaking. After overnight digestion, samples were acidified to pH < 3 with the addition of formic acid (final concentration ∼1% formic acid). Samples were then desalted on C18 StageTips and evaporated in a vacuum concentrator, as previously described (30).

### TMT labeling of peptides

For peptide labeling with TMT (11-plex), desalted peptides were reconstituted in 100 μL of 50 mM HEPES. Each 0.8 mg vial of TMT reagent was reconstituted in 41 μL of anhydrous acetonitrile and added to the corresponding peptide sample for 1 hour at room temperature. The labeling of samples with TMT reagents was completed as shown in Figure 4A. The labeling reactions were quenched with 8 μL of 5% hydroxylamine at room temperature for 15 minutes with shaking, evaporated in a vacuum concentrator, and desalted on C18 StageTips.

The TMT-11 plex labeled samples were fractionated by Strong Cation Exchange (SCX) using StageTips. One SCX StageTip was prepared per sample using 3 plugs of SCX material (3M, cat. #2251) topped with 2 plugs of C18 material. StageTips were conditioned with 100 μL MeOH, 100 μL 80% MeCN/0.5% acetic acid, 100 μL 0.5% acetic acid, 100 μL 0.5% acetic acid/500 mM NH_4_AcO/20% MeCN, followed by another 100 μL 0.5% acetic acid. Dried samples were resuspended in 250 μL 0.5% acetic acid, loaded onto the stage tips, and washed 2x with 100 μL 0.5% acetic acid. Samples were trans-eluted from C18 material onto the SCX with 100 μL 80% MeCN/0.5% acetic acid, and consecutively eluted using 3 buffers with increasing pH. The first elution was with pH 5.15 buffer (50 mM NH_4_AcO/20% MeCN), followed by pH 8.25 buffer (50 mM NH_4_HCO_3_/20% MeCN), and finally with pH 10.3 buffer (0.1% NH_4_OH/20% MeCN). Three eluted fractions were resuspended in 200 μL 0.5% acetic acid (to reduce the concentration of MeCN) and subsequently desalted on C18 stage tips, as previously described.

### Liquid chromatography and mass spectrometry

Desalted peptides were resuspended in 9 μL of 3% MeCN/0.1% FA and analyzed by online nanoflow liquid chromatography tandem mass spectrometry (LC-MS/MS) using a Q Exactive Plus MS (ThermoFisher Scientific) coupled on-line to a Proxeon Easy-nLC 1200 (ThermoFisher Scientific). For each sample, 4 μL was loaded onto a microcapillary column (360 μm outer diameter x 75 μm inner diameter) containing an integrated electrospray emitter tip (10 μm), packed to approximately 24 cm with RepoSil-Pur C18-AQ 1.9 μm beads (Dr. Maisch GmbH) and heated to 50°C. The HPLC solvent A was 3% MeCN/0.1% FA and the solvent B was 90% MeCN/0.1% FA. The SCX fractions were run with a 110-minute method, which used the following gradient profile: (min:%B) 0:2; 1:6; 83:30; 94:60; 95:90; 100:90; 101:50; 110:50 (with the last two steps at a flow rate of 500 nL/minute).

The Q Exactive was operated in the data-dependent mode acquiring HCD MS/MS scans (resolution = 35,000 for TMT11-plex) after each MS1 scan (resolution = 70,000) on the 12 most abundant ions within a 2 second cycle time using an MS1 target of 3 x 10^6^ and an MS2 target of 5 x 10^4^. The maximum ion time utilized for MS/MS scans was 105 milliseconds. The HCD normalized collision energy was set to 31; the dynamic exclusion time was set to 30 seconds, the peptide match was set to “preferred”, and the isotope exclusion functions were enabled; charge exclusion was enabled for charge states that were unassigned, 1, and >6.

### Analysis of mass spectrometry data

Collected data were analyzed using Spectrum Mill software package v6.1pre-release (Agilent Technologies). Nearby MS scans with a similar precursor m/z were merged if they were within ± 60 seconds retention time and ± 1.4 m/z tolerance. MS/MS spectra were excluded from searching if they failed the quality filter by not having a sequence tag length 0 or did not have a precursor MH+ in the range of 750-4000. All extracted spectra were searched against a UniProt database containing human reference proteome sequences. Search parameters included: parent and fragment mass tolerance of 20 p.p.m., 30% minimum matched peak intensity, and ‘calculate reversed database scores’ enabled. The digestion enzyme search parameter used was Trypsin Allow P, which allows K-P and R-P cleavages. The missed cleavage allowance was set to 4. Fixed modifications were carbamidomethylation at cysteine. TMT labeling was required at lysine, but peptide N termini were allowed to be either labeled or unlabeled. Allowed variable modiciations were protein N-terminal acetylation and oxidized methionine. Individual spectra were automatically assigned a confidence score using the Spectrum Mill autovalidation module. Score at the peptide mode was based on a target-decoy false discovery rate (FDR) of 1%. Protein polishing autovalidation was then applied using an auto thresholding strategy. Relative abundances of proteins were determined using TMT reporter ion intensity ratios from each MS/MS spectrum, and the mean ratio is calculated from all MS/MS spectra contributing to a protein subgroup. Proteins identified by ≥2 unique peptides were considered for the dataset.

### Generation of proteomic lists for ERM and OMM

To select cutoffs for proteins biotinylated by TurboID over non-specific binders, proteins were classified as ERM or OMM true positive proteins as established prior (9). Proteins were also classified as mitochondrial matrix proteins (false positive proteins), as determined by GOCC (GO:0005759, but no annotations for IMM, OMM, IMS, or membrane).

To calculate optimal cutoffs, the true positive rate (TPR) and false positive rate (FPR) were calculated at each possible TMT log_2_ ratio. TPR is defined as the fraction of true positive proteins above the TMT ratio in question and FPR is defined as the fraction of false positive proteins above the TMT ratio in question. TMT log_2_ ratios for TurboID targeted to the ERM or OMM over wild-type cells were used. Cutoffs were defined by the maximum (TPR-FPR).

To select cutoffs for proteins biotinylated at the indicated organelle membrane over proteins biotinylated by TurboID targeted to the cytosol, proteins were further classified as cytosolic proteins (false positive proteins), as determined by GOCC (GO:0005829, but no annotations for membrane). TMT log_2_ ratios for TurboID targeted to the ERM or OMM over TurboID targeted to the cytosol were used. To calculate optimal cutoffs, the TPR and FPR were calculated at each possible TMT log_2_ ratio; cutoffs were defined by the maximum (TPR-FPR). After applying both cutoffs to each replicate, the lists were intersected to produce the final proteome lists.

### Generation of proteomic lists for split-TurboID

To select cutoffs for proteins biotinylated by TurboID over non-specific binders, proteins were classified as gold positive ER-mitochondria contact proteins based on prior literature (8, 16, 31, 53–75) or as mitochondrial matrix proteins (false positive proteins), as determined by GOCC (GO:0005759, but no annotations for IMM, OMM, IMS, or membrane). TMT log_2_ ratios for split-TurboID (+R+B) over split-TurboID (+R-B) and for split-TurboID (-R+B) over split-TurboID (-R-B) were used. The FPR was calculated for each possible TMT log_2_ ratio, and the cutoff was determined by setting the false discovery rate (FDR) < 0.1.

To select cutoffs for proteins biotinylated at the ER-mitochondria contacts by split-TurboID over proteins biotinylated by TurboID targeted to the cytosol, proteins were further classified as cytosolic proteins (false positive proteins), as determined by GOCC (GO:0005829, but no annotations for membrane). TMT log_2_ ratios for split-TurboID (+R+B) over TurboID targeted to the cytosol and for split-TurboID (-R+B) over TurboID targeted to the cytosol were used. The FPR was calculated for each possible TMT log_2_ ratio, and the cutoff was determined by setting the false discovery rate (FDR) < 0.1.

After applying both cutoffs to each replicate, the lists were intersected to produce the final proteome lists. FKBP1 and mTOR were removed from the lists as peptides from FKBP and FRB in the introduced split-TurboID constructs would confound these findings. We also generated an additional list intersecting split-TurboID (+R+B) and split-TurboID (-R+B) lists.

### Additional analysis of proteomic data

For specificity analysis, proteomes of each dataset were annotated by gene ontology annotations using AmiGO2. For each list, the proportion of proteins with mitochondrial outer membrane (GO:0005741) and/or endoplasmic reticulum membrane (GO:0005789) annotations was calculated to generate specificity data.

For sensitivity analysis, the proportion of the Gold+ ER-mitochondria contact protein list detected by each proteomic dataset was calculated to generate sensitivity data.

For gene ontology and clustering analysis of the combined proteomic dataset, the final proteome was analyzed using Cytoscape (v3.7.2) (75). A network was generated based on protein-protein interactions from the STRING database using a confidence score cutoff of 0.2 and 0 maximum additional interactors. Clustering was performed using a Markov clustering algorithm with the granularity parameter set to 1.8. Gene ontology analysis was performed for proteins from each cluster using AmiGO2.

### Subcellular fractionation and preparation of MAM fractions

Subcellular fractions were isolated from HEK293T. Cells were harvested after washing with PBS and centrifuged at 600g for 5 minutes. All of the following procedures were performed at 4°C. Cell pellets were resuspended in resuspension buffer 1 (225 mM mannitol, 75 mM sucrose, 0.1 mM EGTA, 30 mM Tris-HCl (pH 7.4)) and homogenized using a Dounce homogenizer. Cell homogenates were centrifuged twice at 600g for 5 minutes to pellet nuclei and intact cells. The supernatants were collected and centrifuged at 7,000g for 10 minutes to pellet crude mitochondria. The pellets were resuspended in resuspension buffer 2 (225 mM mannitol, 75 mM sucrose, 30 mM Tris-HCl (pH 7.4)) and the supernatants were saved for separation of ER and cytosolic fractions. The resuspended crude mitochondrial fractions were then centrifuged at 10,000g for 10 minutes, and the pellets were resuspended in MRB buffer (mitochondria resuspending buffer) (250 mM mannitol, 5 mM HEPES, 0.5 mM EGTA (pH 7.4)). Mitochondria and MAM were separated from the resuspended pellets using percoll medium (225 mM mannitol, 25 mM HEPES, 1 mM EGTA, 30% Percoll (v/v) (pH 7.4)) by ultracentrifugation (SW55-Ti rotor, Optima-XE100 ultracentrifuge, Beckman Coulter) at 95,000g for 30 minutes. The diffused floating band (MAM) and pellets (mitochondria) were collected in separate tubes. Mitochondrial pellets and MAM were diluted in MRB buffer and centrifuged at 6,300g for 10 minutes. Pelleted mitochondria were collected; supernatants from the MAMs were subjected to ultracentrifugation at 100,000g for 60 minutes to pellet the MAM fraction. For ER and cytosolic fractions, the supernatants were centrifuged at 20,000g for 30 minutes, followed by ultracentrifugation at 100,000g for 60 minutes. The pellets (ER) and supernatants (cytosolic) were collected in separate tubes. All of the isolated subcellular fractions were resuspended with protein lysis buffer supplemented with 1x protease inhibitor and phosphatase inhibitor cocktail (ThermoFisher Scientific). Protein concentrations were determined by BCA and 20 μg of protein per sample were separated by SDS-PAGE.

## Competing Financial Interests

The authors declare no competing financial interest.

---

**Supp. Figure 1.**
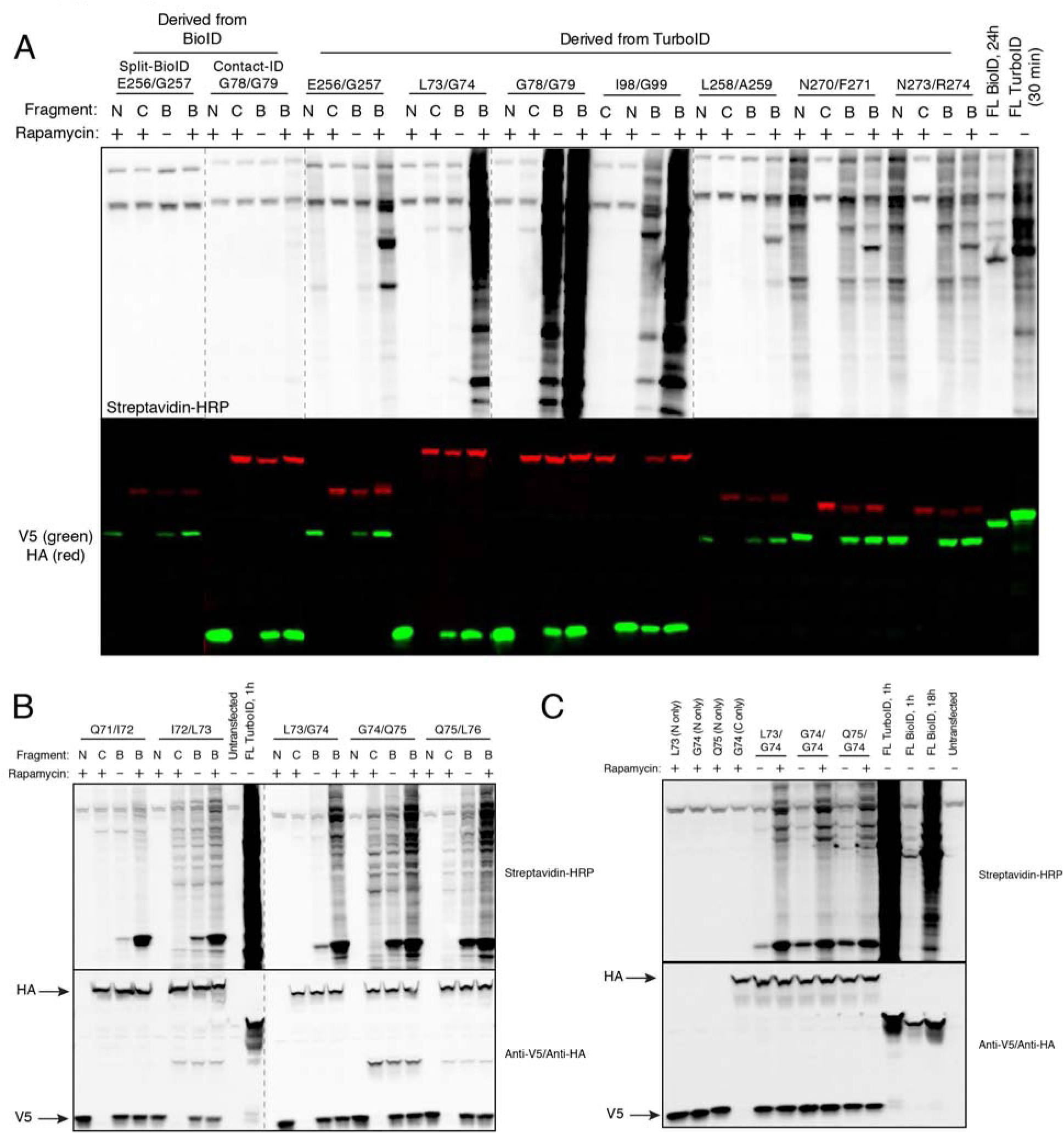
Screening of TurboID split sites. Related to. Figure 1. *(A)* Comparison of split sites in HEK293T cells. Cells transiently transfected with the indicated constructs were treated with 50 μM biotin and 100 nM rapamycin for 24 hours, and whole cell lysates were analyzed by streptavidin blotting. For each construct pair, lanes are shown with both fragments present (B), N-terminal fragment only (N), or C-terminal fragment only (C). Anti-V5 and anti-HA staining show expression levels of N- and C-terminal fragments, respectively. Full-length TurboID (30 minutes labeling) and full-length BioID were included for comparison. Dashed lines indicate separate blots performed at the same time and developed simultaneously. Quantification of biotinylation activity is shown in Figure 1C. *(B)* Testing of additional split sites around the most promising split (73/74) from the first round of screening in HEK293T cells. Cells transiently transfected with the indicated constructs were treated with 50 μM biotin and 100 nM rapamycin for 1 hour, and whole cell lysates were analyzed by streptavidin blotting. For each split site, the N-terminal and C-terminal fragments were either expressed alone (N or C) or both together (B). Anti-V5 and anti-HA staining show expression levels of N- and C-terminal fragments, respectively. Full-length TurboID (treated with 50 μM biotin for 1 hour) and untransfected samples were included for comparison. *(C)* Testing additional combinations of N-terminal and C-terminal fragments from (C) in HEK293T cells. Samples were analyzed as described in (B), with 1 hour of labeling. Full-length TurboID and BioID treated with 50 μM biotin for 1 hour, full-length BioID treated with 50 μM biotin for 18 hours, and untransfected samples were included for comparison.

**Supp. Figure 2.**
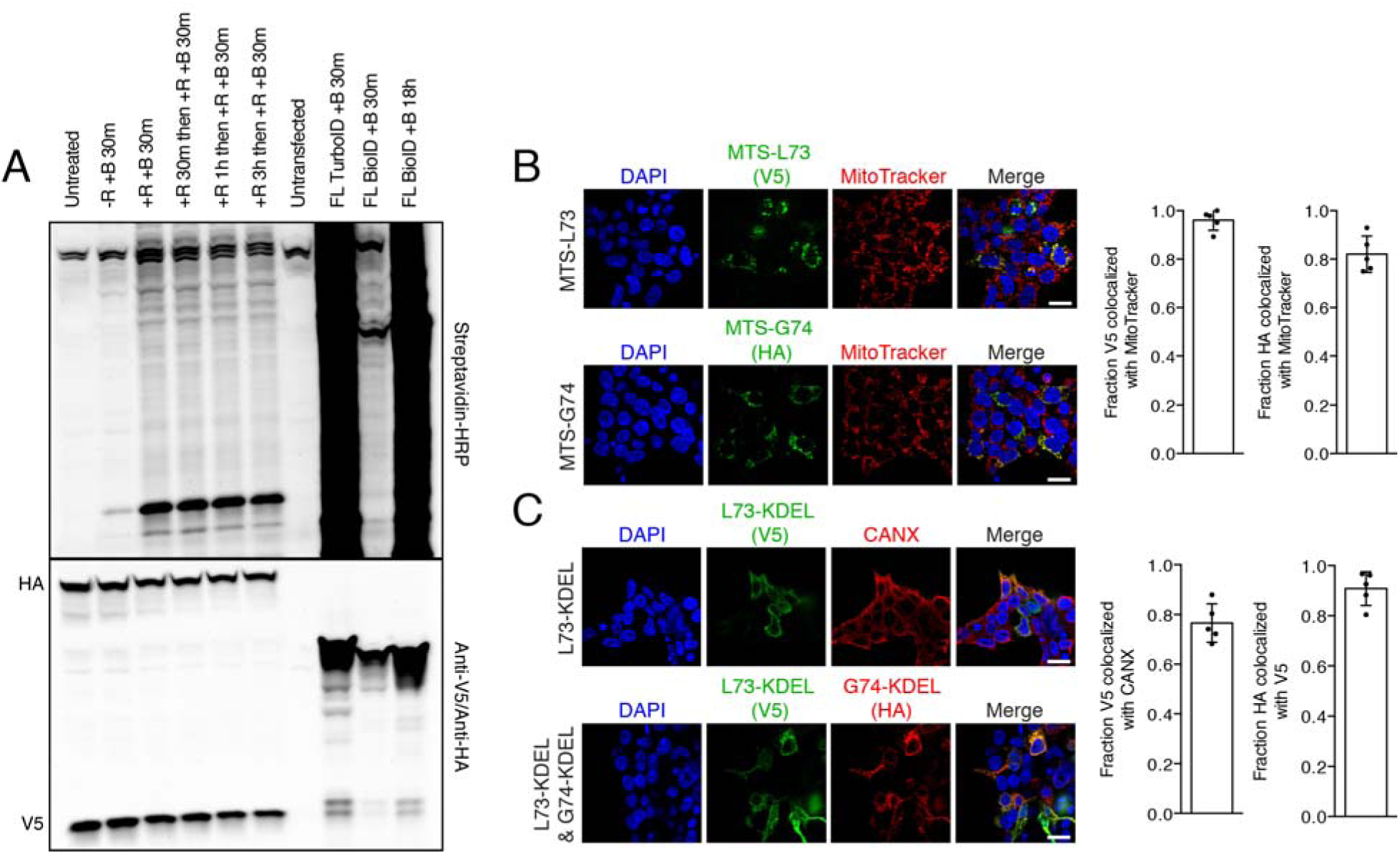
Additional characterization of low-affinity split-TurboID and colocalization of organelle-targeted constructs. Related to. Figure 2. *(A)* Split-TurboID reconstitutes with fast kinetics. Cells transiently transfected with split-TurboID constructs were pre-treated with 100 nM rapamycin for the indicated time before labeling with biotin. Whole cell lysates were analyzed by streptavidin blotting, and anti-V5 and anti-HA staining show expression levels. Full-length TurboID (30 minutes) and full-length BioID (30 minutes and 18 hours) were included for comparison. *(B)* Confocal fluorescence imaging of split-TurboID targeted to the mitochondrial matrix. Cells were labeled with 500 nM MitoTracker for 30 minutes, fixed, and stained with anti-V5 or anti-HA to detect the respective fragment. Scale bars, 20 μm. Colocalization of V5 or HA with MitoTracker is quantified on the right. Quantitation from 5 fields of view per condition are included. *(C)* Confocal fluorescence imaging of split-TurboID (L73/G74) targeted to the ER lumen. Cells transiently transfected with the N-terminal fragment alone were fixed and stained with anti-V5 to detect the N-terminal fragment, and with CANX, which is an ER marker. Cells transiently transfected with both fragments were fixed and stained with anti-V5 and anti-HA to detect the respective fragments simultaneously. Scale bars, 20 μm. Colocalization of either V5 and CANX or V5 and HA is quantified on the right. Quantitation from 5 fields of view per condition are included.

**Supp. Figure 3.**
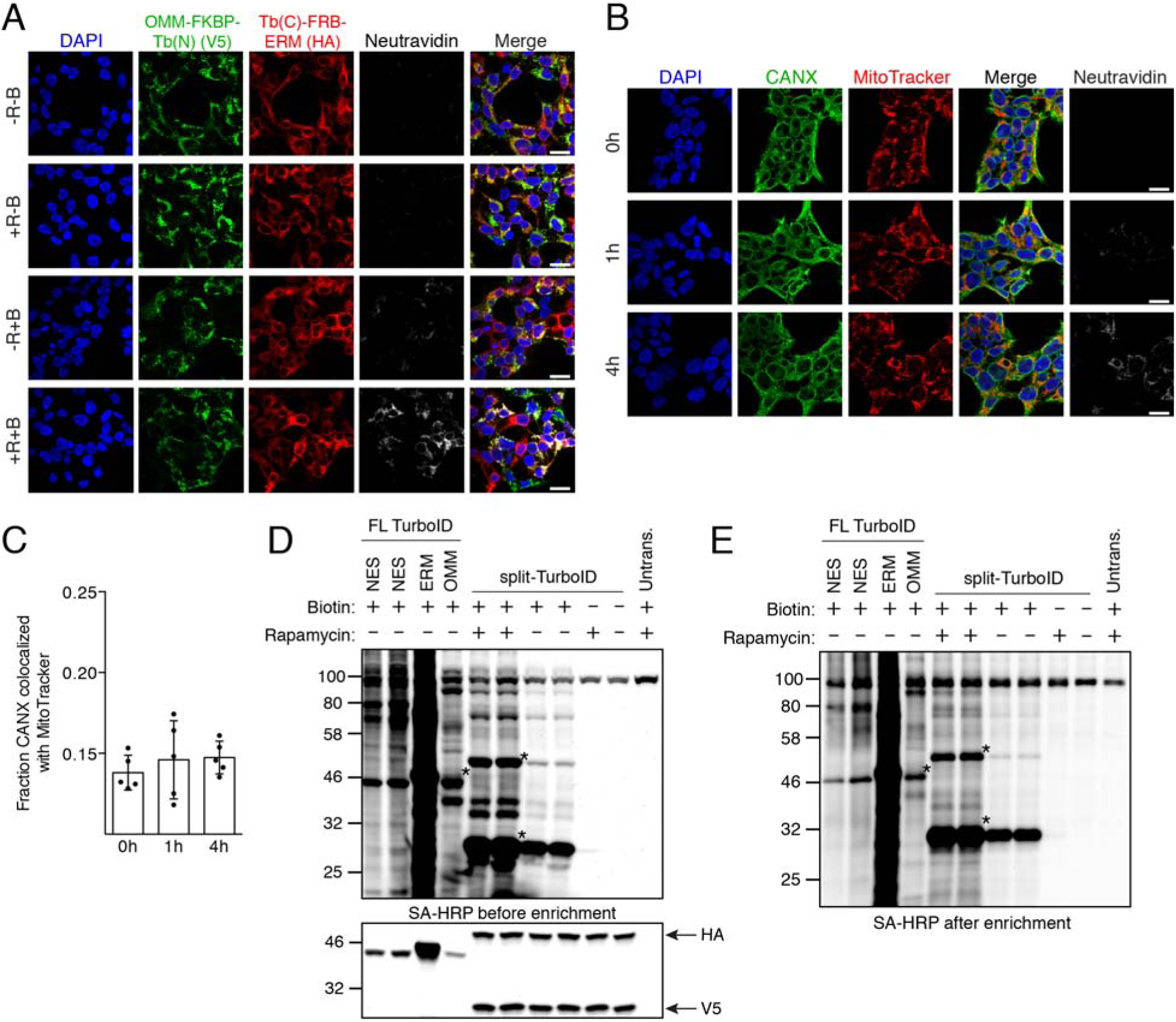
Characterization of stable HEK293T cell lines expressing split-TurboID used for ER-mitochondria proteomics. Related to. Figure 3. *(A)* Confocal fluorescence imaging of split-TurboID targeted to the OMM and ERM. Cells stably expressing split-TurboID constructs were treated with 50 μM biotin and 100 nM rapamycin for 1 hour, then fixed and stained with anti-V5 to detect the OMM-targeted N-terminal fragment (Tb(N)), anti-HA to detect the ERM-targeted C-terminal fragment (Tb(C)), and neutravidin-647 to detect biotinylated proteins. Scale bars, 20 μm. *(B)* Cells stably expressing OMM/ERM-targeted split-TurboID constructs were treated with 50 μM biotin and 100 nM rapamycin for 0, 1, or 4 hours. Cells were labeled with 500 nM MitoTracker for 30 minutes then fixed and stained with anti-CANX, which is an ER marker and neutravidin-647 to detect biotinylated proteins. *(C)* Quantification of colocalization between CANX and MitoTracker in (B) (using a threshold of >15 times background signal). Quantitation from 5 fields of view per condition are included. *(D)* Enrichment of biotinylated proteins for proteomics. HEK293T cells stably expressing OMM/ERM-targeted split-TurboID constructs were treated with 50 μM biotin and 100 nM rapamycin for 4 hours. HEK293T cells stably expressing NES-, OMM-, or ERM-targeted full-length TurboID were treated with 50 μM biotin for 1 minute. Whole cell lysates were analyzed by streptavidin blotting. Asterisks indicate ligase self-biotinylation. *(E)* Samples were generated as in (D). Biotinylated proteins were enriched from lysates using streptavidin beads, eluted, and analyzed by streptavidin blotting. Asterisks indicate ligase self-biotinylation

**Supp. Figure 4.**
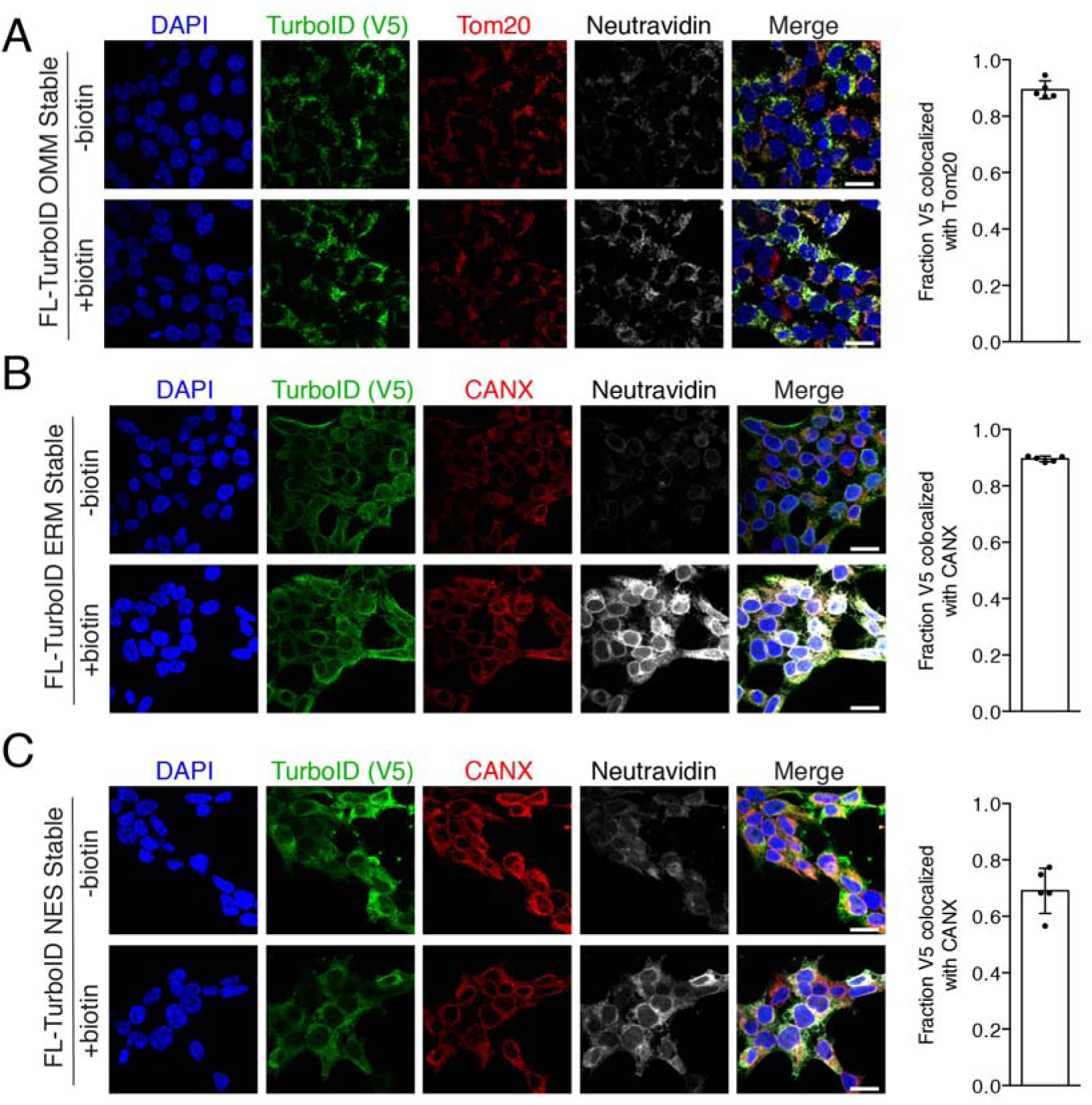
Characterization of stable HEK293T cell lines expressing full-length TurboID used for ER-mitochondria proteomics. Related to Figure 3. *(A)* Confocal fluorescence imaging of full-length TurboID targeted to the OMM. Cells were treated with 50 μM biotin for 10 minutes, then fixed and stained with anti-V5 to detect the OMM-targeted TurboID, Tom20, which is a mitochondrial marker, and neutravidin-647 to detect biotinylated proteins. Scale bars, 20 μm. Colocalization of V5 and Tom20 is quantified on the right. Quantitation from 5 fields of view per condition are included. *(B)* Confocal fluorescence imaging of full-length TurboID targeted to the ERM. Cells were treated with 50 μM biotin for 10 minutes, then fixed and stained with anti-V5 to detect the ERM-targeted TurboID, CANX, which is an ER marker, and neutravidin-647 to detect biotinylated proteins. Scale bars, 20 μm. Colocalization of V5 and CANX is quantified on the right. Quantitation from 5 fields of view per condition are included. *(F)* Confocal fluorescence imaging of full-length TurboID fused to NES for cytosolic targeting. Cells were treated with 50 μM biotin for 10 minutes, then fixed and stained with anti-V5 to detect TurboID, CANX, which is an ER marker, and neutravidin-647 to detect biotinylated proteins. The merged column shows that cytosolic TurboID does not fully overlap and is distinct from the ER marker. Scale bars, 20 μm. Colocalization of V5 and CANX is quantified on the right. Quantitation from 5 fields of view per condition are included.

**Supp. Figure 5.**
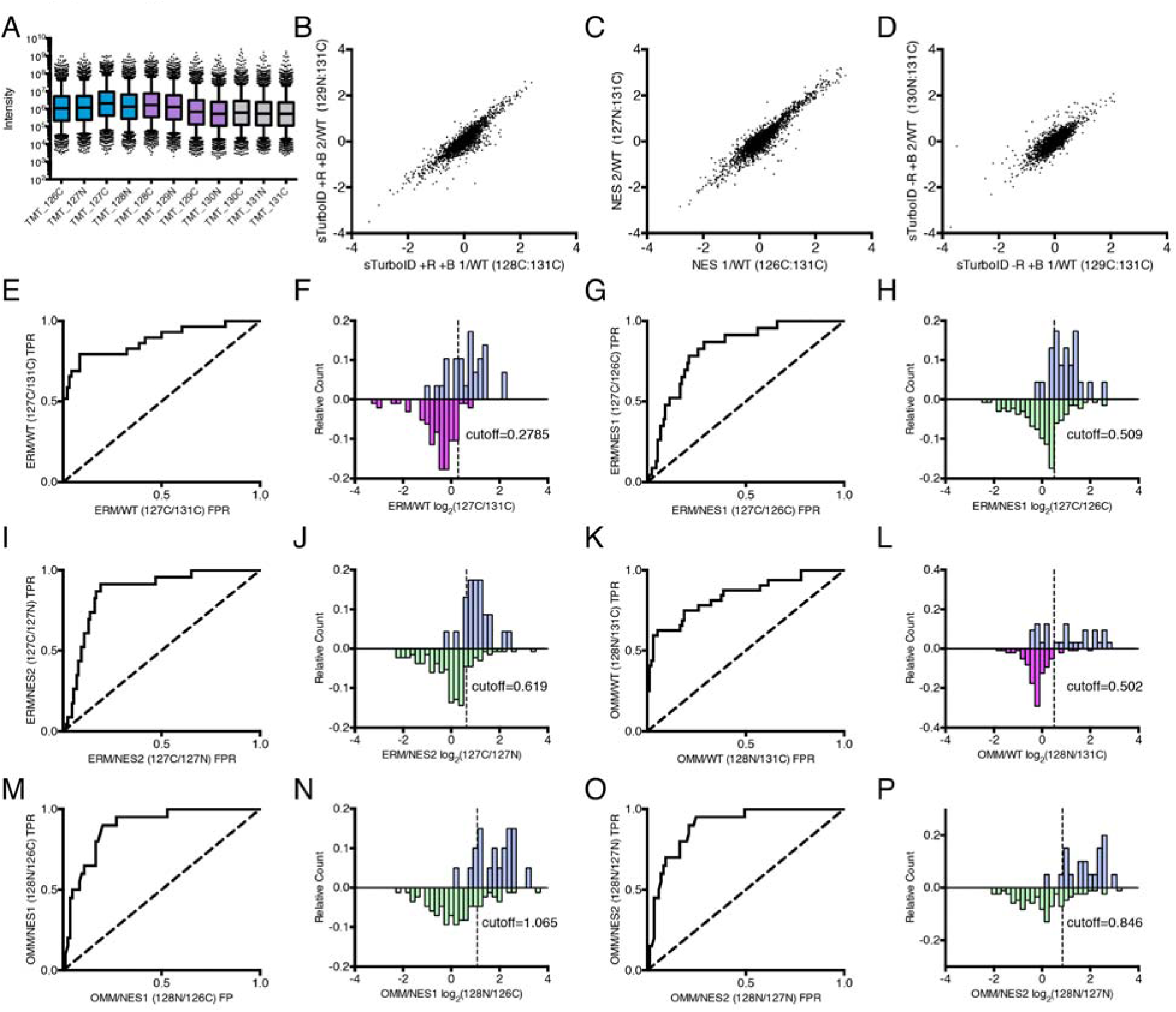
Additional analyses of proteomic data. Related to. Figure 4. *(A)* Intensity plots for TMT-labeled samples showing that sample input is relatively uniform with controls at slightly lower levels, as expected. *(B-D)* Scatterplots of log_2_ ratios for replicates of full-length TurboID-NES (B) (r^2^=0.8712), split-TurboID (+R+B) (C) (r^2^=0.8081), and split-TurboID (-R+B) (D) (r^2^=0.6799). *(E)* ROC (receiver operating characteristic) curve (true positive rate (TPR) versus false positive rate (FPR)) for ERM-targeted TurboID compared to the wild-type sample. *(F)* Histograms of the distributions of true positive proteins (blue, top) and false positive (mitochondrial matrix) proteins (pink, bottom). The cutoff was determined by the maximum (TPR-FPR) and corresponds to FDR < 0.083. *(G)* ROC curve for ERM-targeted TurboID compared to the cytosolic TurboID (replicate 1). *(H)* Histograms of the distributions of true positive proteins and false positive (cytosolic) proteins (green, bottom). The cutoff was determined by the maximum (TPR-FPR) and corresponds to FDR < 0.288. *(I)* ROC curve for ERM-targeted TurboID compared to the cytosolic TurboID (replicate 2). *(J)* Histograms of the distributions of true positive and false positive proteins. The cutoff was determined by the maximum (TPR-FPR) and corresponds to FDR < 0.189. *(K)* ROC curve for OMM-targeted TurboID compared to the wild-type sample. *(L)* Histograms of the distributions of true positive proteins false positive proteins. The cutoff was determined by the maximum (TPR-FPR) and corresponds to FDR < 0.052. *(M)* ROC curve for OMM-targeted TurboID compared to the cytosolic TurboID (replicate 1). *(N)* Histograms of the distributions of true positive proteins and false positive proteins. The cutoff was determined by the maximum (TPR-FPR) and corresponds to FDR < 0.2. *(O)* ROC curve for ERM-targeted TurboID compared to the cytosolic TurboID (replicate 2). *(P)* Histograms of the distributions of true positive proteins and false positive proteins. The cutoff was determined by the maximum (TPR-FPR) and corresponds to FDR < 0.247.

**Supp. Figure 6.**
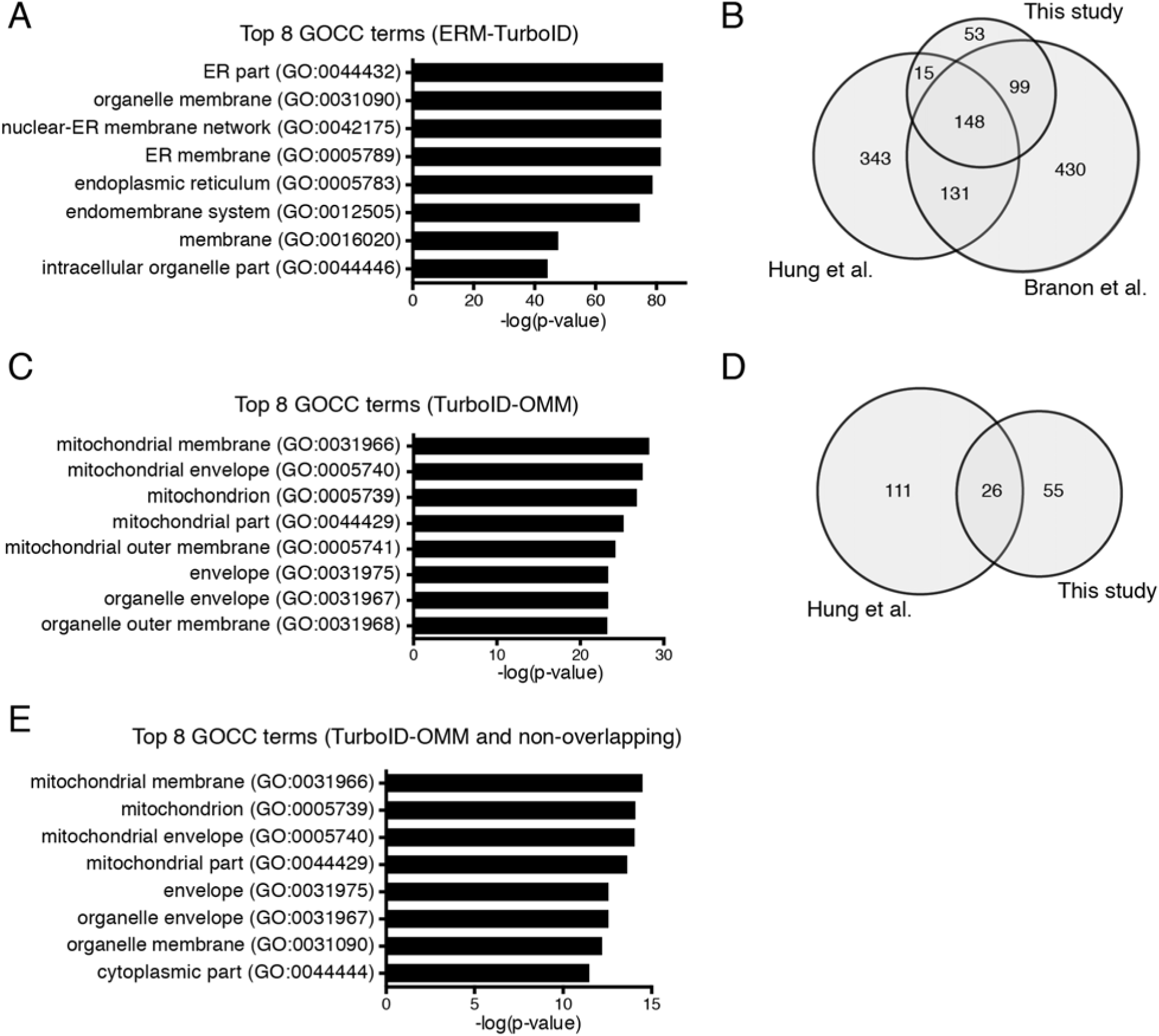
Additional analyses of proteomic data. Related to. Figure 4. *(A)* Top eight GOCC terms (PANTHER) after GO enrichment analysis of the final proteome for ERM-targeted TurboID. *(B)* Venn diagram showing overlap of proteomic lists compared to previously published datasets using TurboID and APEX (7, 8). *(B)* Top eight GOCC terms (PANTHER) after GO enrichment analysis of the final proteome for OMM-targeted TurboID. *(C)* Venn diagram showing overlap of proteomic lists compared to previously published dataset using APEX (2). *(D)* Top eight GOCC terms (PANTHER) after GO enrichment analysis of the final proteome for OMM-targeted TurboID that does not overlap with the previously published dataset from Hung et al (8).

**Supp. Figure 7.**
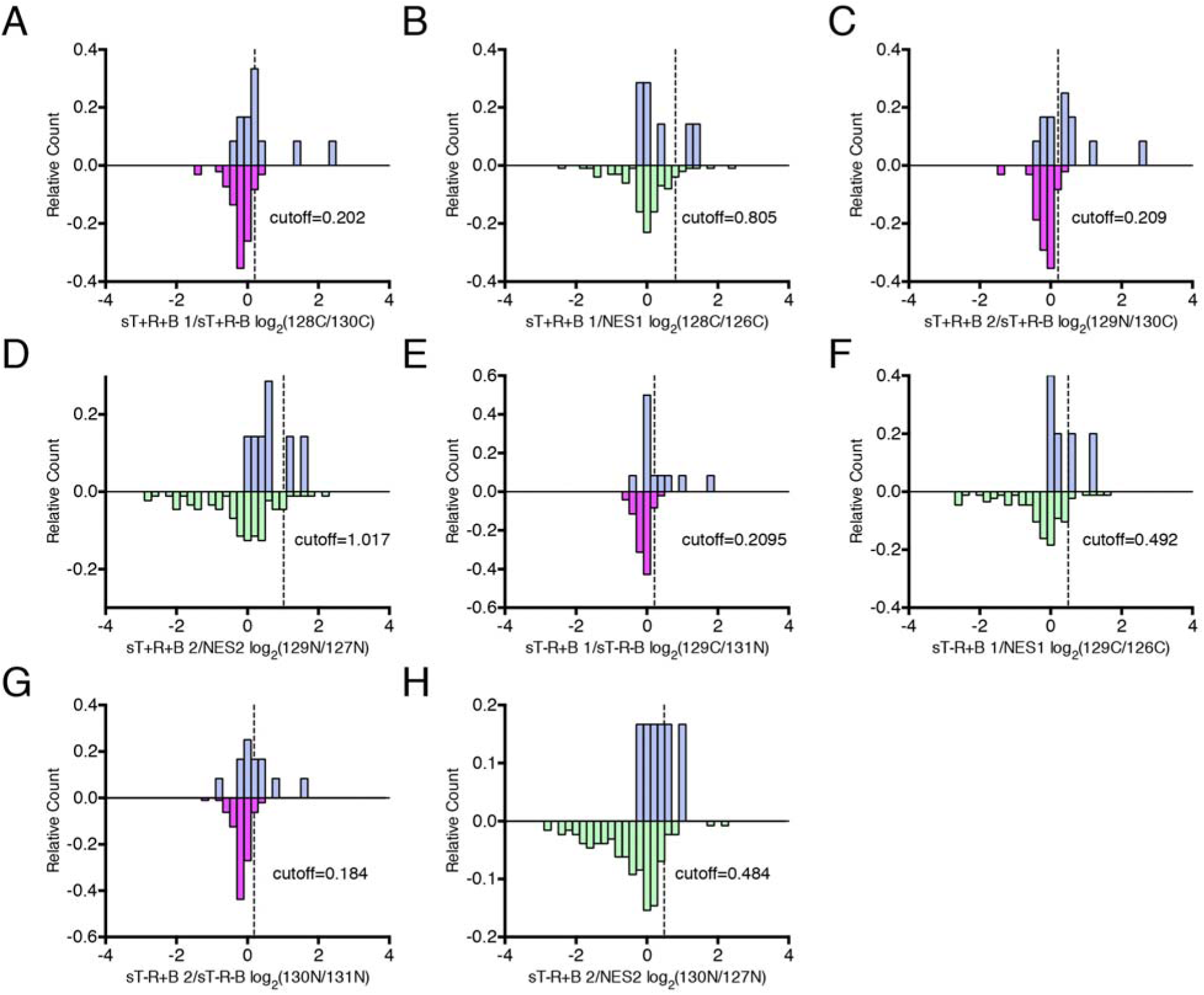
Histograms used to filter split-TurboID proteomic data. Related to. Figure 4 **and** Figure 5. *(A)* Histograms of the distributions of true positive proteins (blue, top) and false positive (mitochondrial matrix) proteins (pink, bottom) for split-TurboID (+R+B) (replicate 1). *(B)* Histograms of the distributions of true positive proteins (blue, top) and false positive (cytosolic) proteins (green, bottom) for split-TurboID (+R+B) (replicate 1). *(C)* Histograms of the distributions of true positive proteins and false positive proteins for split-TurboID (+R+B) (replicate 2). *(D)* Histograms of the distributions of true positive proteins and false positive proteins for split-TurboID (+R+B) (replicate 2). *(E)* Histograms of the distributions of true positive proteins and false positive proteins for split-TurboID (-R+B) (replicate 1). *(F)* Histograms of the distributions of true positive proteins and false positive proteins for split-TurboID (-R+B) (replicate 1). *(G)* Histograms of the distributions of true positive proteins and false positive proteins for split-TurboID (-R+B) (replicate 2). *(H)* Histograms of the distributions of true positive proteins and false positive proteins for split-TurboID (-R+B) (replicate 2). Cutoffs were determined by FDR < 0.1 and are represented in the graphs as dashed lines.

**Supp. Figure 8.**
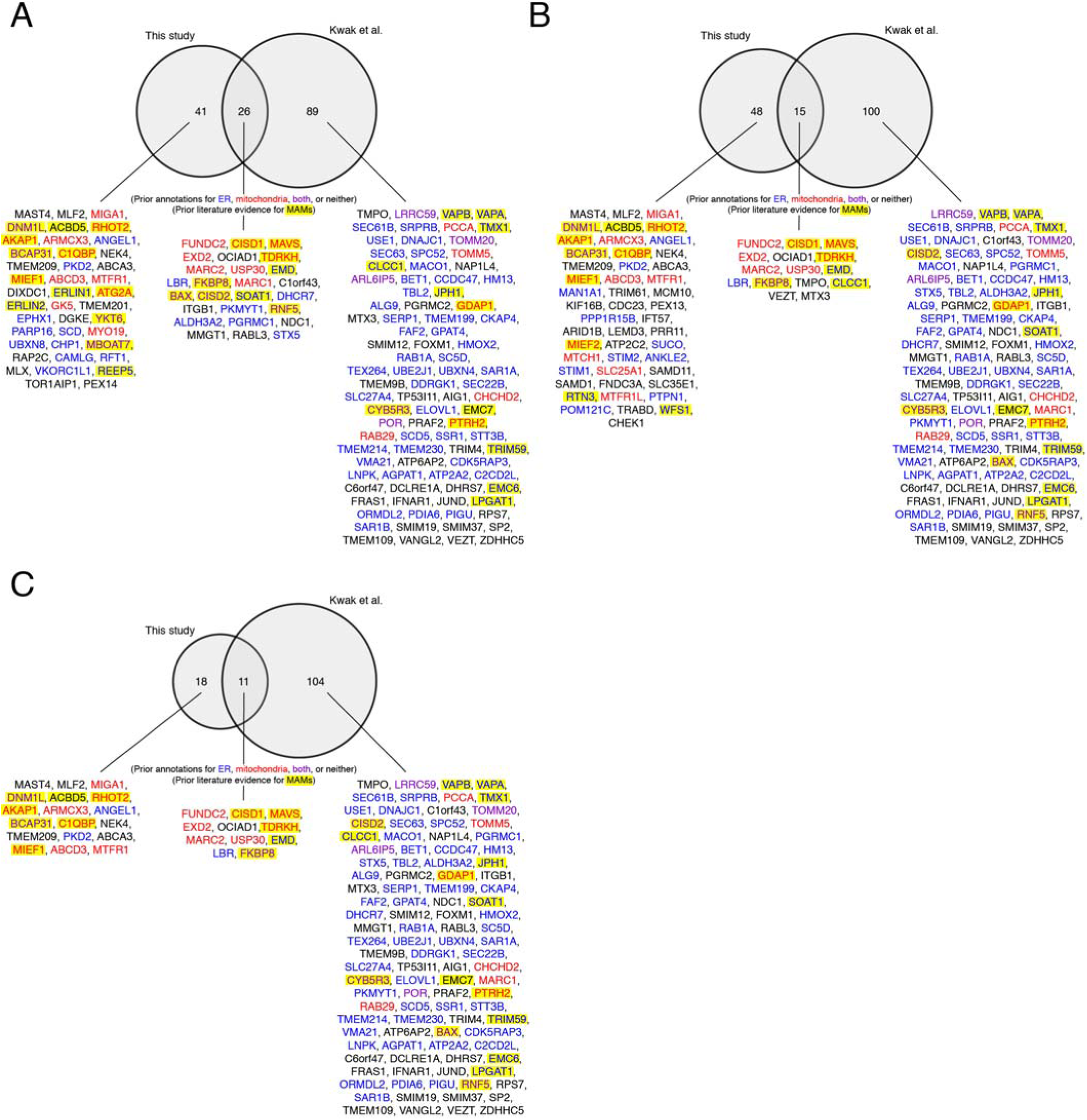
Comparison of our split-TurboID ER-mitochondria proteomes with Contact-ID ER-mitochondria proteome (Kwak et al.) (21). Venn diagram comparing proteome lists using split-TurboID (+rapamycin in (A), -rapamycin in (B), and +rapamycin/-rapamycin overlap in (C) to contact-ID proteome list (Kwak et al (21)). Proteins that were previously annotated with ER, mitochondria, both, or neither were labeled blue, red, purple, or black, respectively. Proteins with previous literature evidence for association with MAMs (15-17, 52-101) are highlighted in yellow.

**Supp. Figure 9.**
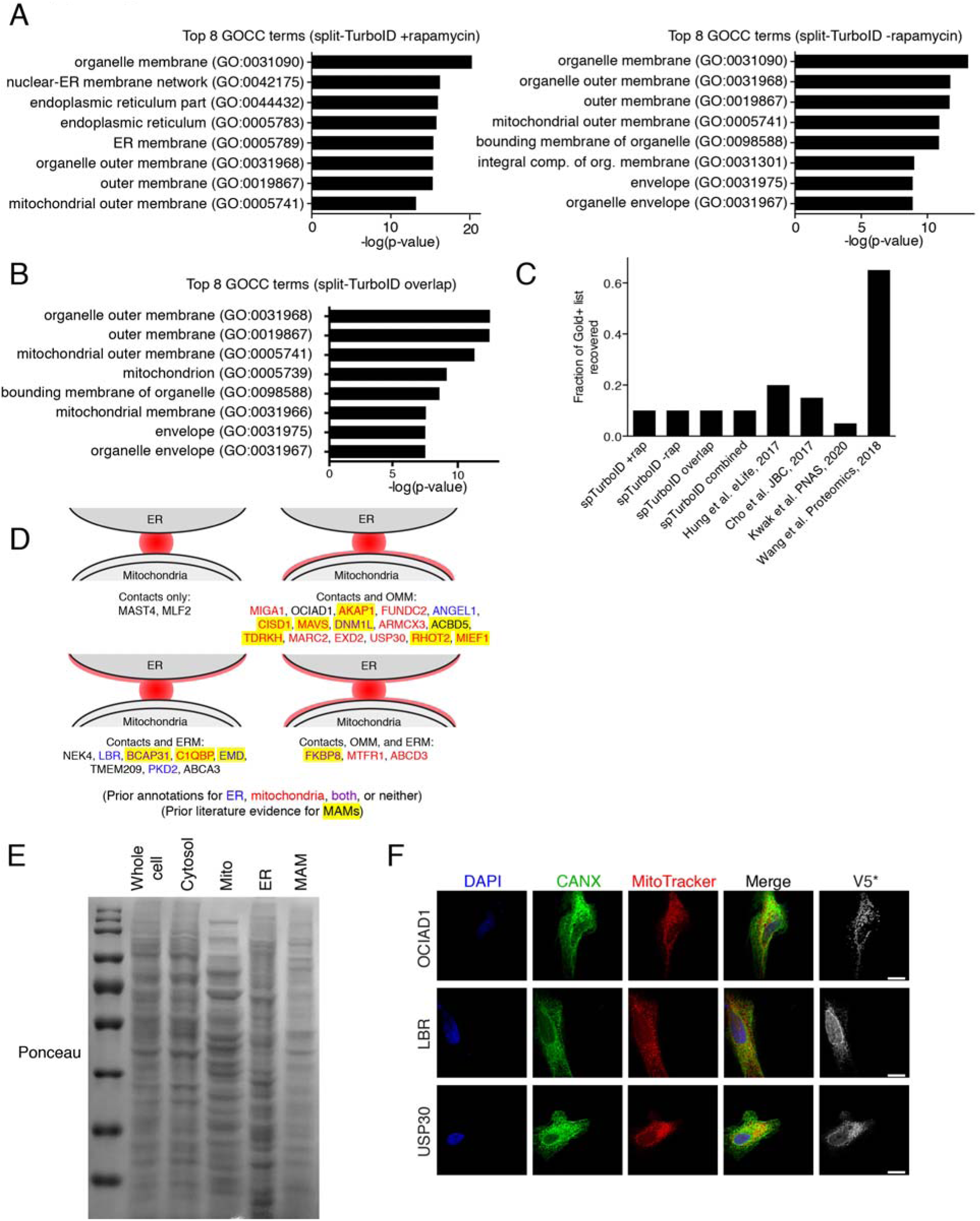
Analysis and validation of proteomic hits. Related to Figure 5 and Figure 6. *(A)* Top eight GOCC terms (PANTHER) for the split-TurboID +rapamycin proteome (67 proteins) (left) and -rapamycin proteome (63 proteins) (right). *(B)* Top eight GOCC terms (PANTHER) after GO enrichment analysis of +rapamycin/-rapamycin overlap proteome. *(C)* Sensitivity analysis for proteomic datasets generated using split-TurboID compared to previous datasets studying ER-mitochondria contacts or MAMs. Bar graphs show the percentage of proteins in our manually curated true positive (“Gold+”) list detected in each proteome. *(D)* Distinct localization patterns for ER-mitochondria contact resident proteins. For the proteins enriched in both +rapamycin and -rapamycin split-TurboID datasets, some were enriched exclusively by split-TurboID (top left), while others were enriched by split-TurboID *and* TurboID-OMM (top right), suggesting dual localization to contacts and OMM. Proteins enriched at contacts and ERM shown at bottom left, and proteins enriched in three locations shown at bottom right. Gene names are colored by prior GOCC or literature annotation. *(E)* Representative Ponceau stain for Western blotting of proteins in samples generated from subcellular fractionation in Figure 6A. *(E)* Confocal fluorescence imaging of candidate ER-mitochondria contact proteins OCIAD1, LBR, and USP30 overexpressed in HeLa cells. Constructs were introduced by transient transfection. Cells were incubated with 500 nM MitoTracker for 30 minutes, then fixed and stained with anti-CANX to detect the ER and anti-V5 to detect the gene of interest. V5 signals are not normalized across the different constructs tested. Quantification of colocalization between CANX and MitoTracker is shown in Figure 6C.

